# Cell-type aware regulatory landscapes governing monoterpene indole alkaloid biosynthesis in the medicinal plant *Catharanthus roseus*

**DOI:** 10.1101/2024.04.23.590703

**Authors:** Chenxin Li, Maite Colinas, Joshua C. Wood, Brieanne Vaillancourt, John P. Hamilton, Sophia L. Jones, Lorenzo Caputi, Sarah E. O’Connor, C. Robin Buell

**Affiliations:** Center for Applied Genetic Technologies, University of Georgia, Athens, GA, USA; Department of Crop and Soil Sciences, University of Georgia, Athens, GA, USA; Department of Natural Product Biosynthesis, Max Planck Institute for Chemical Ecology, Jena, Germany; Institute of Plant Breeding, Genetics, and Genomics, University of Georgia, Athens, Georgia, USA

## Abstract

In plants, the biosynthetic pathways of some specialized metabolites are partitioned into specialized or rare cell types, as exemplified by the monoterpenoid indole alkaloid (MIA) pathway of *Catharanthus roseus* (Madagascar Periwinkle), the source of the anti-cancer compounds vinblastine and vincristine. In the leaf, the *C. roseus* MIA biosynthetic pathway is partitioned into three cell types with the final known steps of the pathway expressed in the rare cell type termed idioblast. How cell-type specificity of MIA biosynthesis is achieved is poorly understood. Here, we generated single-cell multi-omics data from *C. roseus* leaves. Integrating gene expression and chromatin accessibility profiles across single cells, as well as transcription factor (TF) binding site profiles, we constructed a cell-type-aware gene regulatory network for MIA biosynthesis. We showcased cell-type-specific transcription factors as well as cell-type-specific *cis*-regulatory elements. Using motif enrichment analysis, co-expression across cell types, and functional validation approaches, we discovered a novel idioblast specific TF (<u>Id</u>ioblast <u>M</u>YB1, CrIDM1) that activates expression of late stage vinca alkaloid biosynthetic genes in the idioblast. These analyses not only led to the discovery of the first documented cell-type-specific TF that regulates the expression of two idioblast specific biosynthetic genes within an idioblast metabolic regulon, but also provides insights into cell-type-specific metabolic regulation.

## Introduction

An emerging feature of plant specialized metabolism is the spatial and temporal restriction of biosynthetic gene expression ^1^, some of which are confined to rare and specialized cells within an organ ^2^. The medicinal plant *Catharanthus roseus* produces monoterpene indole alkaloids (MIAs), including vinblastine and vincristine (also known as vinca alkaloids) that are clinically used to treat various cancers ^3^. The MIA biosynthetic pathway can be conceptually divided into four stages: the methyl erythritol phosphate (MEP) pathway that provides the precursor to the monoterpene moiety of MIAs, the iridoid stage that generates the monoterpene moiety of MIAs, the alkaloid scaffolding stage, and finally the late alkaloid stage that further decorates MIA (Supplementary Table 1). The MIA pathway genes in *C. roseus* display intricate cell-type specific expression patterns. The MEP and iridoid stages of the pathway are exclusively expressed in a specialized vasculature associated cell type, the inner phloem associated parenchyma (IPAP) ^4–6^. The alkaloid scaffolding steps are expressed in the epidermis ^4,5,7^, and the final known steps of the pathway are restricted to a rare and specialized cell type termed idioblast ^7,8^, which are scattered throughout the leaf ^9,10^. In addition to its economic importance as the source of chemotherapeutic medications, the intricate partitioning of the MIA pathway into multiple cell types highlights *C. roseus* as a model system for investigating cell-type specific regulation of plant specialized metabolism.

Several transcription factors (TFs) have been identified as regulators of the MIA biosynthetic pathway in *C. roseus* ^11–17,17–20^, primarily in the context of jasmonate (JA)-induction of this pathway. Major known regulators of the MIA pathway include MYC2 ^14,20^, bHLH iridoid synthesis (BIS) family TFs ^15,18,19^, and Octadecanoid-derivative Responsive Catharanthus AP2-domain (ORCA) family TFs ^12,13,17,21^, all of which mediate JA induction of the MIA pathway. However, since all currently available studies on transcriptional regulation of the MIA pathway have relied on whole organ (bulk) samples, how the pathway is regulated at the cell type level remains unknown. Furthermore, MYC2, BIS, and ORCA families TFs have been shown to activate the pathway up to the alkaloid scaffolding stage of the pathway, and to date, no cell-type-specific regulators for the late-stage portion of the pathway have been identified.

Here, we apply single cell multiome (gene expression and accessible chromatin profiles from the same nucleus) to investigate the regulatory landscapes of the MIA biosynthetic pathway in mature *C. roseus* leaves at the cell type level. Using co-expression across single cells, transcription factor binding site (TFBS) profiles, and cell-type-aware TFBS accessibility, we constructed a knowledge-based gene regulatory network (GRN) for this biosynthetic pathway. Our analyses uncovered a new idioblast specific MYB TF that through functional genomics approaches, we showed regulates the expression of two idioblast specific biosynthetic genes which are co-regulated within an idioblast metabolic regulon. This study discovered a new regulatory component pertinent to the final steps of vinblastine and vincristine biosynthesis in *C. roseus* and furthers our understanding of cell-type-specific regulation of plant specialized metabolism.

## Results

### 1. The cell-type specific expression patterns of MIA biosynthetic genes are reflected in single cell multiome profiles

To investigate the regulation of MIA biosynthetic genes (Supplementary Table 1, Supplementary Fig. 1A) at the single cell resolution, we generated dual gene expression and chromatin accessibility profiles across single cells. We first isolated intact nuclei (Supplementary Fig. 1B-I) from mature *C. roseus* leaves and constructed replicated single cell multiome (RNA-seq and assay for transposase accessible chromatin followed by sequencing [ATAC-seq]) libraries using the 10x Genomics Multiome Kit (Supplementary Table 2). For gene expression, we obtained gene expression profiles for a total of 8,803 high quality nuclei and 18,532 expressed genes (Fig. 1A, Supplementary Fig. 2, Supplementary Table 3).

**Fig. 1.**
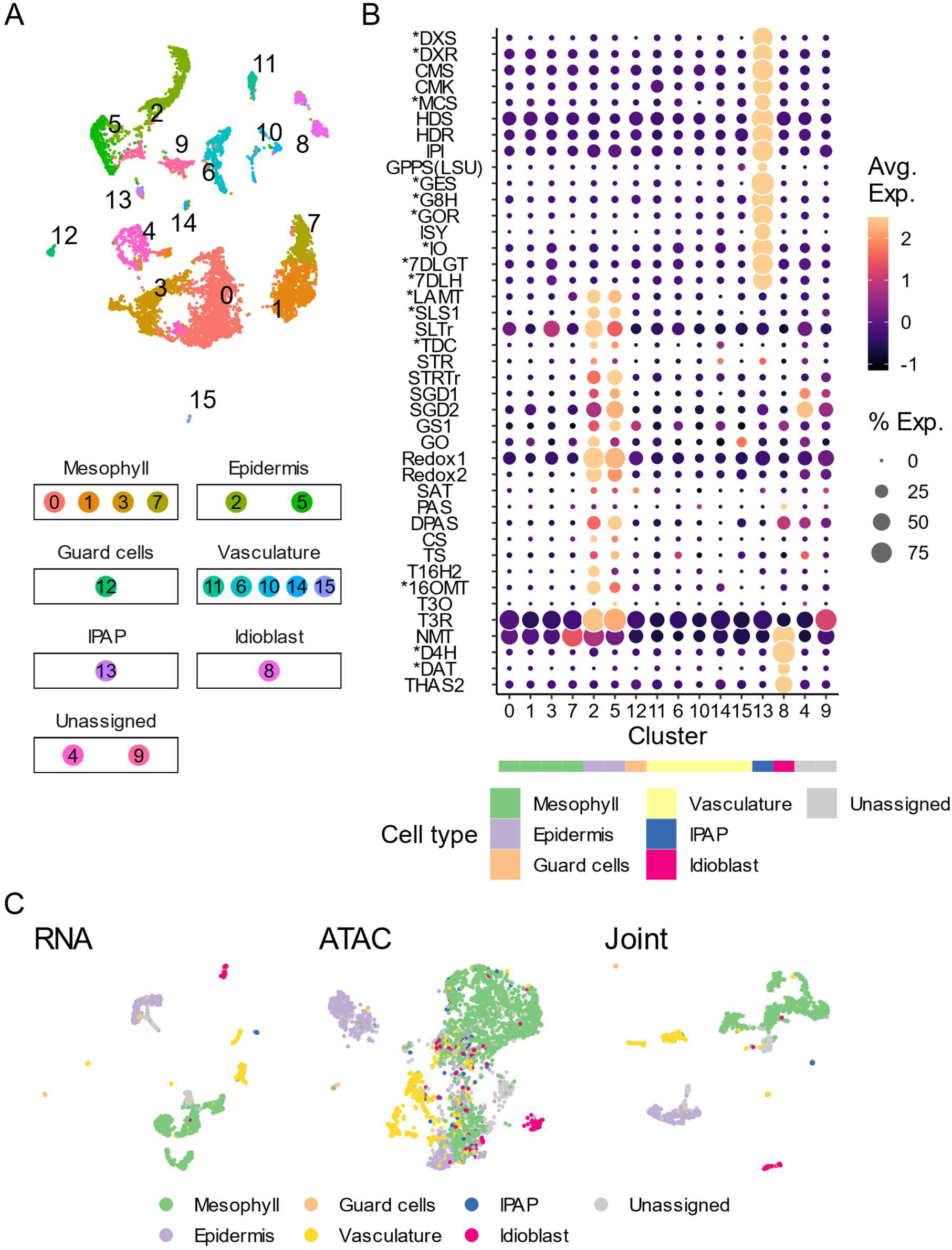
Cell-type specific expression of MIA biosynthetic genes is recapitulated in a leaf single cell multiome dataset. A. UMAP of nuclei containing high-quality RNA-seq data (*n* = 8,803), color coded by cell clusters. B. Gene expression heatmap of MIA biosynthetic genes across cell clusters detected in (A). Rows are biosynthetic genes and transporters, which are ordered from upstream to downstream (see also Supplementary Table 1). Asterisks denote matching cell type specificity with previously reported RNA *in situ* hybridization results ^4–7^. Color scale shows the average scaled expression of each gene at each cell cluster. Dot size indicates the percentage of cells where a given gene is detected. The predicted cell type for each cell cluster is annotated by the color strip below the x-axis (see also Supplementary Fig. 3B and Supplementary Table 5). C. UMAP of nuclei containing both high-quality RNA-seq and ATAC-seq data (*n* = 3,542 nuclei for all three UMAP), color coded by cell types. From left to right: UMAP based on gene expression assay, chromatin accessibility assay, and joint analysis.

We first examined the gene expression data of this multiome dataset (Fig. 1A). Cell clustering patterns are highly similar across the three biological replicates (Supplementary Fig. 3A). Using previously established marker genes ^4–8^ (Supplementary Table 4), we identified major cell types of leaf (e.g., mesophyll, epidermis, and vasculature) as well as two rare cell types in which MIA biosynthetic genes were expressed (i.e., IPAP and idioblast) (Supplementary Fig. 3B). Mesophyll and epidermis were the most abundant cell types, accounting for 54% and 18% of assayed nuclei, respectively. Consistent with their rare nature, IPAP and idioblast accounted for only 1% and 4% of assayed nuclei, respectively (Supplementary Fig. 3C). We found that the MIA biosynthetic pathway was organized into three discrete cell types (Fig. 1B). The MEP and iridoid stages (up to 7-DLH, Supplementary Fig. 1A) of the pathway were exclusively expressed in the IPAP cells. The following stage, which includes most of the alkaloid steps, was expressed in the epidermis. Finally, the last four known steps of the pathway were only expressed in the idioblast. The data were highly consistent with recently published single cell RNA-seq results using protoplasts ^8,22^ and were fully supported by previously reported RNA *in situ* hybridization results (marked with asterisk) ^4–7^.

We next proceeded to analyze chromatin accessibility data to investigate how biosynthetic genes might be regulated to generate cell-type-specific expression patterns. For the chromatin accessibility assay, high quality ATAC-seq nuclei have fraction of fragments in peaks > 0.25, greater than 2,000 ATAC fragments per nuclei, and greater than 1,000 peaks per cell (Supplementary Table 4), resulting in accessibility profiles for a total of 3,765 high quality nuclei and 43,630 accessible chromatin peaks (Fig. 1C). We performed a joint analysis by matching the cell barcodes from both assays. Matching 8,803 high quality nuclei from the RNA-seq assay with 3,765 high quality nuclei for the ATAC-seq assay, the joint analysis resulted in an intersecting set of 3,542 nuclei containing both high-quality RNA-seq and ATAC-seq data (Fig. 1C).

### 2. A gene regulatory network for MIA biosynthetic genes integrating co-expression, chromosome accessibility, and transcription factor binding site (TFBS) profiles

To investigate the regulation of MIA biosynthetic genes, we examined chromatin accessibility landscapes across cell types. ATAC-seq fragments were highly enriched at transcription start and end sites (Supplementary Fig. 4). Among the three biological replicates, 45.9%, 40.9%, and 41.3% of ATAC-seq fragments overlapped transcriptional start sites (Supplementary Fig. 4, Supplementary Table 5). We then defined ATAC-seq peaks using MACS2 ^23^; among biological replicates, 48.9%, 48.5% and 48.8% of fragments are within ATAC-seq peaks (Supplementary Fig. 5, Supplementary Table 5). The median length of ATAC-seq peaks was 566-bp (Supplementary Fig. 6A). The chromatin accessibility landscapes were complemented with transcription factor binding site (TFBS) profiles of ORCA3, a well-known master regulator of MIA biosynthesis ^17^, and its tandemly duplicated paralog ORCA4 ^21^ (Fig. 2A, B, Supplementary Table 6). We determined TFBS profiles for ORCA3/4 using DNA affinity purification sequencing (DAP-seq) ^24^. Average DAP-seq peak lengths were similar (∼300-bp) between ORCA3 and ORCA4 (Supplementary Fig. 6B, C) with ∼10% of DAP-seq peaks intersecting with ATAC-seq peaks (Supplementary Fig. 6D), consistent with the *in vitro* nature of the DAP-seq assay ^24^. Signal-to-noise ratios at DAP-seq peaks were strong (Supplementary Fig. 6E, F, Supplementary Table 6), comparable to the most high-quality DAP-seq datasets that have been published ^24^.

**Fig. 2.**
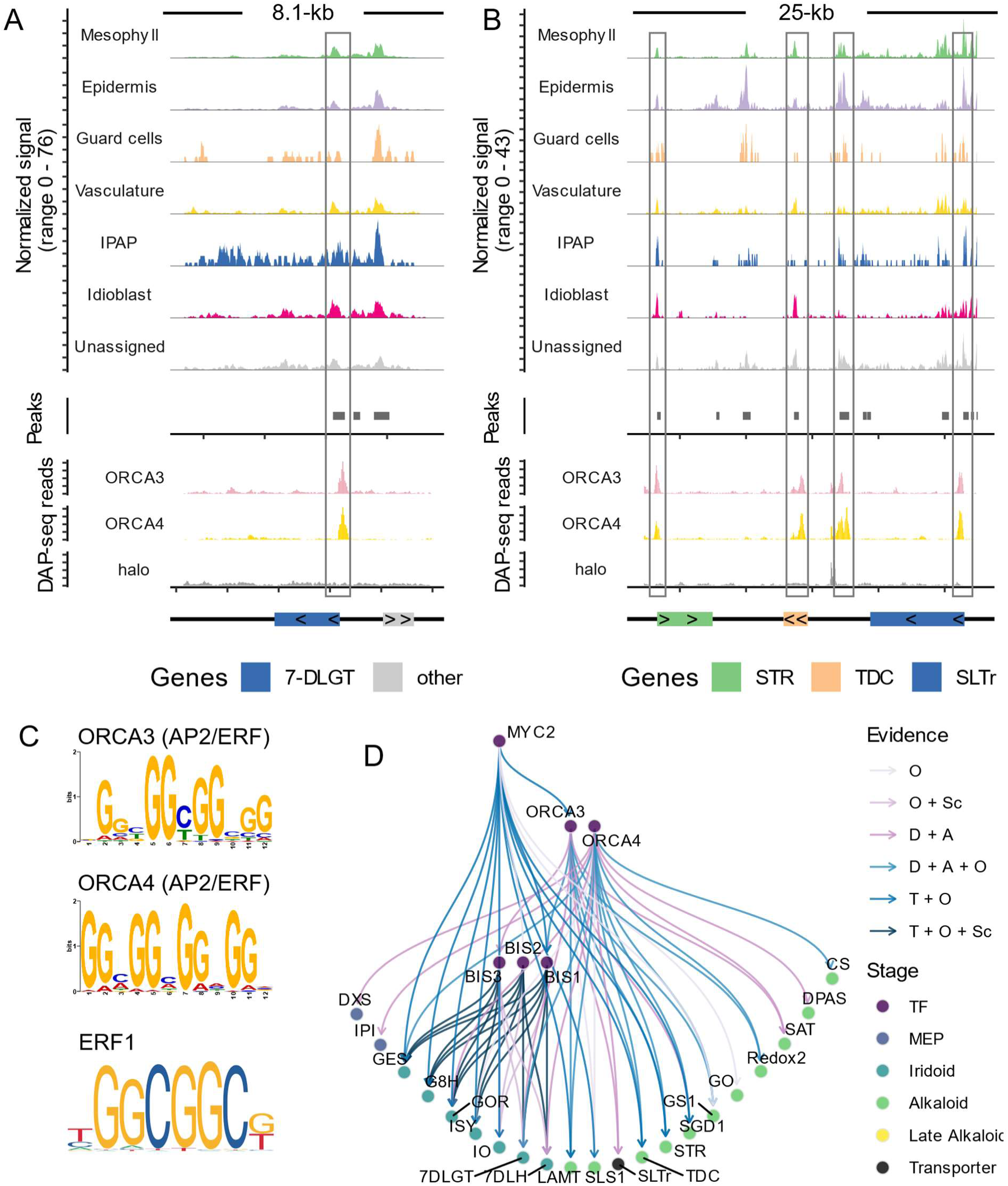
A gene regulatory network for MIA biosynthetic genes integrating chromosome accessibility landscapes and transcription factor binding site profiles. A-B. Coverage plot showing ATAC-seq (upper panels) and DAP-seq (lower panels) signals at the 7-DLGT locus (A) and STR-TCD-SLTr biosynthetic gene cluster (B). Grey boxes highlight DAP-seq peaks that overlap with ATAC-seq peaks. Bottom track indicates the location and length of genes, where the direction of carets (> or <) indicates the strand of a gene. Halo: control DAP-seq experiment using the halo tag (affinity tag) alone. C. DNA motifs enriched in ORCA3/4 binding sites, as well as a reference GCC box/ERF motif^27^. D. A GRN integrating multiple modules of omics data and experimental data. Each node is a gene, color coded by the stage of the biosynthetic pathway. Each edge represents a regulatory relationship, color coded by the type of evidence supporting it. O: upregulated when the TF is overexpressed; Sc: co-expressed across single cells; D: overlapping or within 2-kb to a DAP-seq peak; A: DAP-seq peak accessible; T: promoter activated in a transactivation assay. Gene abbreviations are listed in Supplementary Table 1.

ORCA TFs are known to activate both the alkaloid steps of the biosynthetic pathway (e.g., Strictosidine Synthase [STR] and Tryptophan Decarboxylase [TDC]) ^13,17^ and the upstream iridoid steps (e.g., 7-DLGT) ^21^. 7-DLGT is exclusively expressed in the IPAP cells (Fig. 1B) ^8^ and consistent with its expression specificity, the 7-DLGT locus has a unique chromatin accessibility signal in IPAP cells at both 5’ and 3’ ends of the gene (Fig. 2A). Strong DAP-seq peaks were observed for both ORCA3 and ORCA4 at the 7-DLGT locus, but not for the affinity-tag control (Fig. 2A). These DAP-seq peaks also overlapped with an ATAC-seq peak that was accessible across all cell types. Together with previously reported data that overexpression of ORCA3 or ORCA4 led to the upregulation of 7-DLGT ^21^, 7-DLGT is a direct target of both ORCA3 and ORCA4.

ORCA3 has been reported to bind to the promoters of STR and TDC and activate their expression ^17,21^. STR and TDC are physically clustered on chromosome 3, along with the secologanin transporter SLTr ^8^. Multiple ATAC-seq peaks were detected within this 25-kb biosynthetic gene cluster, all of which were accessible across multiple cell types (Fig. 2B). ORCA3 and ORCA4 displayed similar binding profiles at this biosynthetic gene cluster. Each TF binds a total of four DAP-seq peaks in this region. Consistent with STR and TDC being direct targets of ORCA3, DAP-seq peaks were detected in the promoters of both STR and TDC. ORCA4 has also been reported to activate both STR and TDC in overexpression assays ^21^, and the presence of ORCA4 binding sites suggests ORCA4 can directly activate both STR and TDC. Lastly, since binding sites for ORCA3/4 were detected at the promoter of Secologanin Transporter (SLTr), and as SLTr is highly co-expressed with STR and TDC in the epidermis (Fig. 1B) ^8^, SLTr is likely a direct target for ORCA3/4 as well.

We performed *de novo* motif discovery ^25^ to identify the DNA binding motifs of ORCA3/4. We found that the GCC box motif was enriched among ORCA3/4 binding sites (Fig. 2C). The same GCC box motif was detected regardless of whether we used all DAP-seq peaks as input or only accessible DAP-seq peaks as input. The GCC box is recognized by ethylene responsive factors (ERFs) (Fig. 2C) ^26^ consistent with ORCA family TFs being within the broader AP2/ERF family.

Integrating gene co-expression across single cells, TF binding sites, binding site chromatin accessibility, as well as previously reported overexpression ^14,21^ and reporter transactivation data ^13,15,18,19^, we generated a knowledge-based gene regulatory network (GRN) for the MIA biosynthetic pathway (Fig. 2D). We first queried the expression patterns of previously studied TFs (Supplementary Table 7) ^11–20,28,29^ in our single cell dataset and found that only ORCA4 and BIS1/2/3 displayed cell type specific expression patterns relevant to iridoid and alkaloid biosynthetic genes (Supplementary Fig. 7A). BIS1/2/3 were expressed specifically in the IPAP cells, highly concordant with the iridoid biosynthetic genes that they regulate (Fig. 1B). ORCA4, but not ORCA3, was expressed specifically in the epidermis, albeit only in a small fraction of cells. Thus, ORCA4, but not ORCA3, may contribute to the epidermal specific expression of alkaloid biosynthetic genes such as STR, TDC, and SLTr (Fig. 2B). All other TFs reported in the literature to be associated with MIA biosynthesis were expressed broadly across cell types, or were not expressed in IPAP, epidermis, or idioblast cells (Supplementary Fig 7A).

Based on their co-expression with target genes at the cell type level, BIS1/2/3 and ORCA4 were selected as TF nodes for the GRN. Co-overexpression of MYC2 and ORCA3 was previously reported to strongly activate the iridoid and alkaloid stages of the pathway ^14^, and thus MYC2 and ORCA3 were also included in this network (Fig. 2D). The gene regulatory network contains 66 edges (Supplementary Table 8), which were decorated by the types of evidence: 1) activated by overexpression of TF, 2) co-expressed at the single cell level, 3) overlapping or within 2-kb of a DAP-seq peak, 4) DAP-seq peak accessible, and 5) promoter activated in a transactivation assay (Fig. 2D). We found that the combined actions of MYC2, ORCA3/4, and BIS1/2/3 activate a large section of the MIA pathway, up to the biosynthetic gene encoding Catharanthine Synthase. Evidence also supported multiple feed-forward regulatory loops, where an upstream TF activates both downstream TFs and biosynthetic genes. The downstream TFs in turn activate the same target biosynthetic genes. For example, ORCA3/4 activates iridoid and alkaloid biosynthetic genes, as well as BIS TFs that in turn activate iridoid biosynthetic genes. However, we also found that no regulatory relationships were detected beyond Catharanthine Synthase for these six TFs, consistent with previous reports where overexpression of MYC2, ORCA, and/or BIS TFs led to an increase in early-stage alkaloid metabolites (e.g., strictosidine), but not late-stage alkaloid such as vinblastine ^14,21^. These observations prompted us to investigate components involved in the regulation of the late MIA pathway.

### 3. Cell-type specific accessible chromatin regions mark late-stage MIA biosynthetic genes

MIA biosynthetic pathway genes downstream of Catharanthine Synthase are sequentially expressed in epidermis (TS, T16H2, 16OMT, T3O, and T3R) and then in idioblast cells (NMT, D4H, DAT, and THAS2) (Fig. 1B, Supplementary Table 1) ^7,8^. T16H2 and 16OMT are consecutive steps of the late MIA pathway, expressed exclusively in the epidermis (Fig. 1B), and physically linked as a biosynthetic gene cluster (Fig. 3A), between which there is another gene encoding a cytochrome P450 that was not expressed in the leaf. At the T16H2/16OMT locus, there are four ATAC-seq peaks. All but one of the peaks were preferentially accessible in the epidermis, consistent with the cell-type specific expression of this gene pair (Fig. 3A). DAT, one of the final known steps of the MIA pathway, is only expressed in the idioblast (Fig. 1B), and its promoter was also specifically accessible in the idioblast (Fig. 3B).

**Fig. 3.**
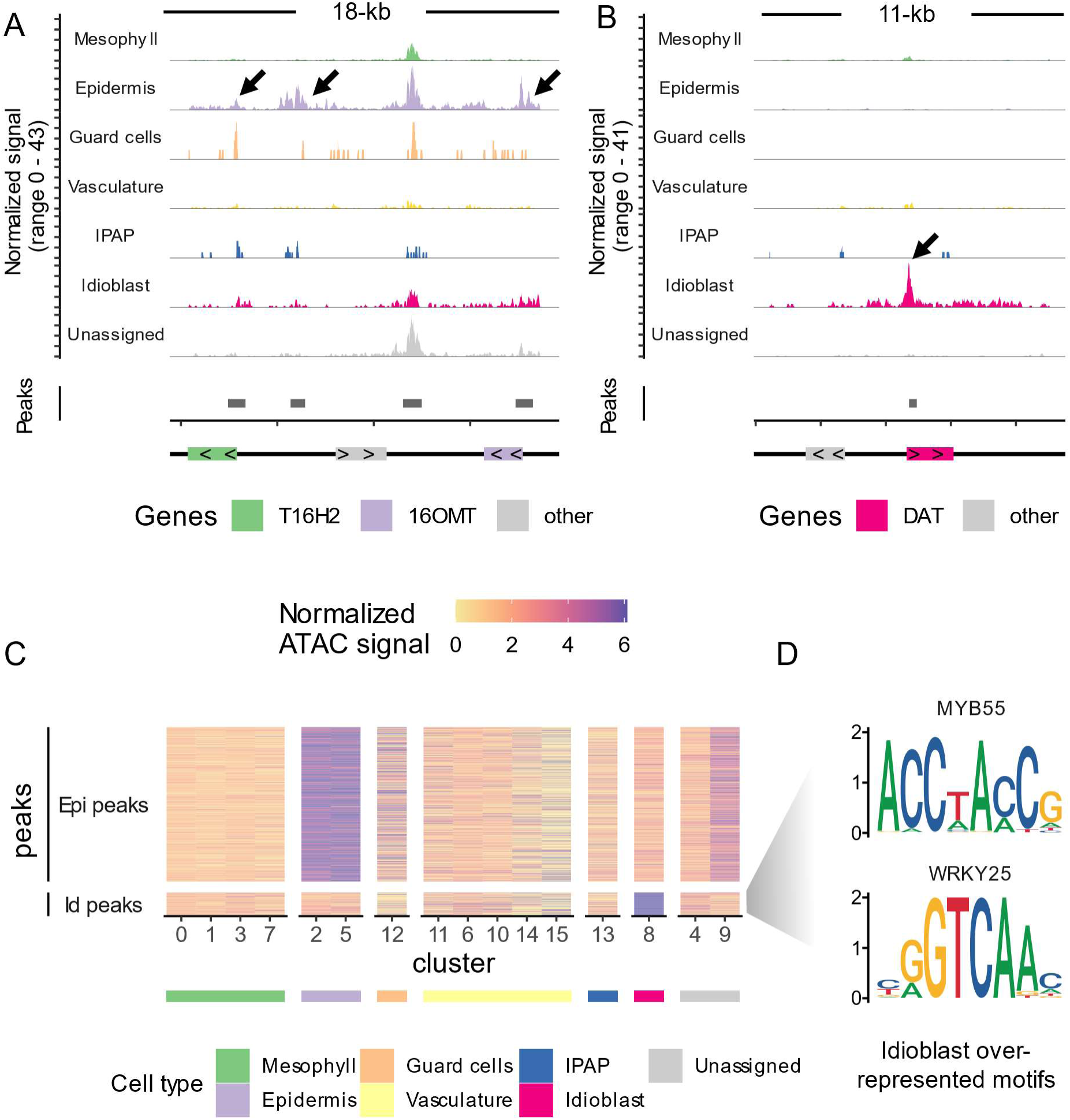
Cell-type specific accessible chromatin regions mark late-stage MIA biosynthetic genes. A-B. Coverage plot showing ATAC-seq signals at the T16H2-16OMT gene pair (A) and DAT locus (B). Arrows highlight cell-type specific ATAC-seq peaks. Bottom track indicates the location and length of genes, where the direction of carets (> or <) indicates the strand of a gene. Grey boxes along the “Peaks” track represent ATAC-seq peaks. C. Heat map showing accessibility of epidermis (Epi) and idioblast (Id) ATAC-seq marker peaks across cell clusters. Each row is an ATAC-seq peak (see also Supplementary Table 9). Each column is a cell cluster. Color scale is maxed out at 90^th^ percentile of normalized ATAC-seq signal. The predicted cell type for each cell cluster is annotated by the color strip below the x-axis (see also Supplementary Fig. 3B). D. TF binding motifs overrepresented among idioblast marker peaks. For motifs overrepresented among epidermis marker peaks, see Supplementary Fig. 7B.

To identify novel regulators for late-stage MIA biosynthetic genes downstream of Catharanthine Synthase, we first detected epidermis and idioblast marker peaks, which are ATAC-seq peaks preferentially accessible in the epidermis or idioblast, but not in any other cell types (Fig. 3C, Supplementary Table 9). We detected 1,050 epidermis-marker peaks and 163 idioblast marker peaks. We next performed a motif enrichment analysis on epidermis marker peaks against the JASPAR plant TF binding motif collection ^27^. We found that homeodomain, ERF, and MYB motifs were overrepresented among epidermis marker peaks (Supplementary Fig. 7B). Homeodomain (e.g., ANTHOCYANINLESS2/ANL2 ^30^), AP2/ERF (e.g., WAX INDUCER1 ^31^), and MYB TFs (e.g., WEREWOLF ^32^) have been reported to control metabolic and developmental processes such as anthocyanin biosynthesis, cuticle development, and trichome development, respectively. Enrichment of these motifs suggests that additional TFs in the homeodomain, ERF, or MYB families may play a role in the regulation of MIA biosynthesis in the epidermis. We also performed motif enrichment analysis on idioblast marker peaks and found that MYB and WRKY type motifs were overrepresented (Fig. 3C), for which we followed up with additional analyses and experiments.

### 4. Candidate WRKY and MYB TFs specifically expressed in the idioblast discovered by gene co-expression analysis

To further understand gene regulation in idioblast cells, we focused our attention on potential metabolic regulators in the idioblast. We performed gene co-expression analysis across cell clusters using graph-based clustering ^33^ and detected tightly co-expressed modules (Fig. 4A). We queried co-expression modules containing MIA biosynthetic genes and detected a single co-expression module for epidermis, IPAP, and idioblast, respectively (Supplementary Table 10). For example, SLS1, which was specifically expressed in the epidermis, was a member of the epidermis co-expression module, whereas the final known steps of the pathway, namely NMT, D4H, DAT, and THAS2 were all members of the idioblast co-expression module (Fig. 4A, Supplementary Table 10). The partitioning of MIA biosynthetic genes into three distinct co-expression modules is similar to a co-expression network constructed from single cell RNA-seq data generated from protoplasts ^8^.

**Fig. 4.**
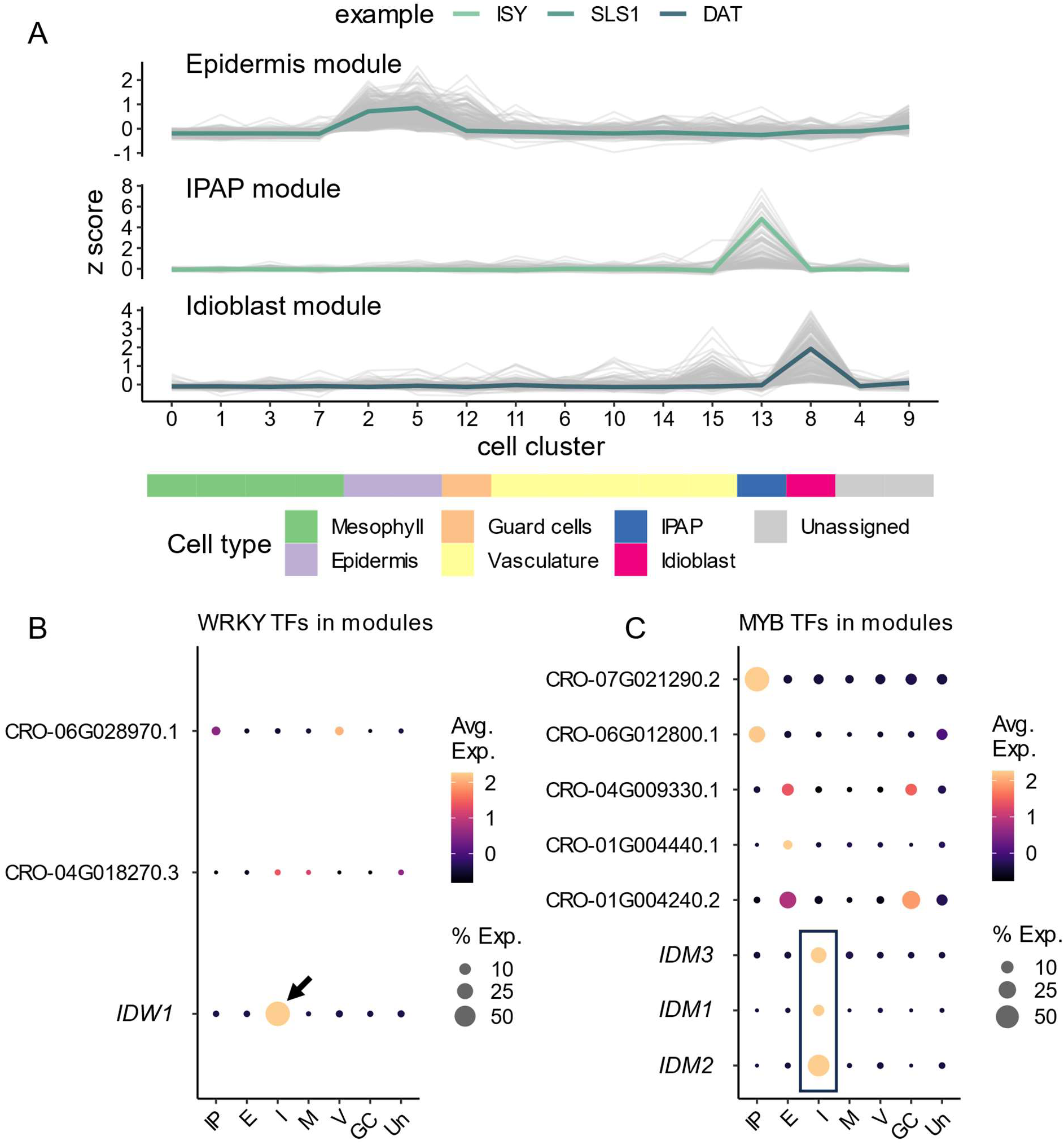
Gene co-expression analysis across cell clusters discovered candidate WRKY and MYB TFs specifically expressed in the idioblast. A. Line graphs showing expression patterns of genes in the epidermis, IPAP, and idioblast co-expression modules (Supplementary Table 10). Grey lines are individual genes, and colored lines are exemplary biosynthetic genes in each module. The predicted cell type for each cell cluster is annotated by the color strip below the x-axis (see also Supplementary Fig. 3B). B-C. Gene expression heatmap of WRKY TFs (B) and MYB TFs (C) across cell types. Color scales show the average scaled expression of each gene for each cell type. Dot size indicates the percentage of cells where a given gene is detected in each cell type. Only WRKY and MYB TFs detected in epidermis, IPAP, or idioblast co-expression modules are presented. Arrow indicates a single WRKY candidate (IDW1: CRO_03G000120) specifically expressed in the idioblats. Box highlights three MYB candidates (IDM1: CRO_05G006800, IDM2: CRO_04G033370, IDM3: CRO_07G002170) specifically expressed in the idioblast.

Since WRKY and MYB motifs were overrepresented among idioblast ATAC-seq marker peaks, we queried WRKY and MYB family TFs within the gene co-expression modules. We identified a single WRKY TF (Fig. 4B) as well as three strong candidates of R2-R3 MYB TFs (Fig. 4C) that were exclusively expressed in the idioblast. We named these candidates *<u>Id</u>ioblast <u>W</u>RKY1* (*IDW1*) and *<u>Id</u>ioblast <u>M</u>YB1/2/3* (*IDM1/2/3*), respectively. All four candidates were induced by a methyl-jasmonate treatment ^34^ (Supplementary Fig. 7C), among which *IDM1* displayed the highest level of induction (log_2_FC = 5.4, or 42-fold increase over control). Since the entire vinblastine biosynthetic pathway is elicited by methyl-jasmonate ^17,18^, the MeJA-responsiveness displayed by these TF candidates suggests they might be transcriptional activators of the pathway. A recent study applied fluorescence activated cell sorting to enrich for idioblast cells prior to RNA-seq ^35^. Consistent with their idioblast specificity, all four TF candidates were detected at high levels in the idioblast fraction of sorted cells, but not in the mesophyll fraction (Supplementary Fig. 7D).

To investigate the phylogenetic relationship among the three MYB candidates, we performed genome-wide identification of MYB domain proteins ^36^ in the *C. roseus* genome ^8^ and detected 92 MYB domain proteins (Supplementary Fig. 8 and Supplementary Fig. 9). We aligned the MYB domains from MYB TFs to produce a phylogeny that includes *C. roseus*, the model species *Arabidopsis thaliana*, *Solanum lycopersicum* (tomato), and *Solanum tuberosum* (potato) MYBs (Supplementary Fig. 8). Tomato and potato MYBs were included to distinguish Solanaceae-specific MYBs against Asterids-specific (encompassing Apocynaceae species that include *C. roseus* and Solanaceae species) MYBs. We found that the three *IDM* candidates were not closely related to each other (Supplementary Fig. 9). Their MYB domains are more similar to MYB TFs in other species than to each other, although they share the same expression pattern. Notably, IDM1 is outgroup to a clade that contains multiple Arabidopsis MYBs that belong to two subclades. One subclade contained MYBs that control trichome and root hair development (MYB0 (GLABRA 1), MYB23, and MYB66 (WEREWOLF)) ^30,37,38^, whereas the other subclade is involved in the regulation of anthocyanin biosynthesis (MYB113/114, MYB75, and MYB90) ^39,40^. IDM2 is outgroup to a clade that contains two less well-characterized Arabidopsis MYBs, MYB6 and MYB8 ^41^. Lastly, IMD3, along with two other *C. roseus* MYBs, is sister to a clade containing Arabidopsis MYB123 (TRANSPARENT TESTA 2/TT2) ^42^, which is involved in proanthocyanidin biosynthesis in the Arabidopsis seed coat (Supplementary Fig. 9).

### 5. IDM1 directly activates the expression of D4H and DAT

To test the functions of IDW1 and IDM1/2/3, we performed overexpression assays followed by RNA-seq to investigate whether overexpression of these TFs affect the expression of the MIA biosynthetic pathway. Coding sequences of *IDW1* and *IDM1/2/3* were cloned downstream of the 35S promoter. The overexpression vectors were transformed into *Agrobacterium tumefaciens* and infiltrated into *C. roseus* petals. In our experience, *C. roseus* petals are much more amendable to agrobacterium-mediated transient expression than leaves, and a highly efficient protocol has been established for petals ^21^. For these reasons, petals were used for transient overexpression assays, instead of leaves. We used GUS as a negative control, as infiltrating agrobacterium affects the expression of the pathway. As a positive control, an engineered MYC2 TF ^14^ and ORCA3 ^17^ were co-infiltrated which have been previously shown to strongly activate the MIA pathway ^14^. The MYC2 coding sequence was previously engineered to carry the D126N mutation, such that it could no longer be post-translationally repressed by the JAZ repressor protein. A combined overexpression treatment of IDW1 and IDM1/2/3 was also tested; a total of seven treatments including controls were assayed.

We performed infiltrations at two agrobacterium titers, 0.1 optical density (OD) and 0.4 OD which is the highest titer that can be used without resulting in wilting of the petals after infiltration (see also Methods). Using triplicated overexpression treatments at 0.1 OD (Supplementary Fig. 10A, Supplementary Table 2), we found that the MYC2-ORCA3 positive control strongly activates the MIA pathway up to the DPAS step (Fig. 5A), consistent with previous reports that these known regulators do not activate later-stage biosynthetic genes downstream of Catharanthine Synthase (Fig. 2D) ^14^. We discovered that one of the MYB candidates, IDM1, activated the expression of both D4H and DAT (Fig. 5A), resulting in a 2.8-fold and 1.68-fold increase in expression relative to the GUS control, respectively. All other overexpression treatments, including the combination of all candidates, did not activate the pathway relative to the GUS control (Fig. 5A). Encouraged by the initial result for IDM1, we examined gene expression profiles at 0.4 OD (Supplementary Fig. 10B, Supplementary Table 2). To control for batch-to-batch variation between experiments, independent GUS controls were included across both 0.1 OD and 0.4 OD experiments (Fig. 5A).

**Fig. 5.**
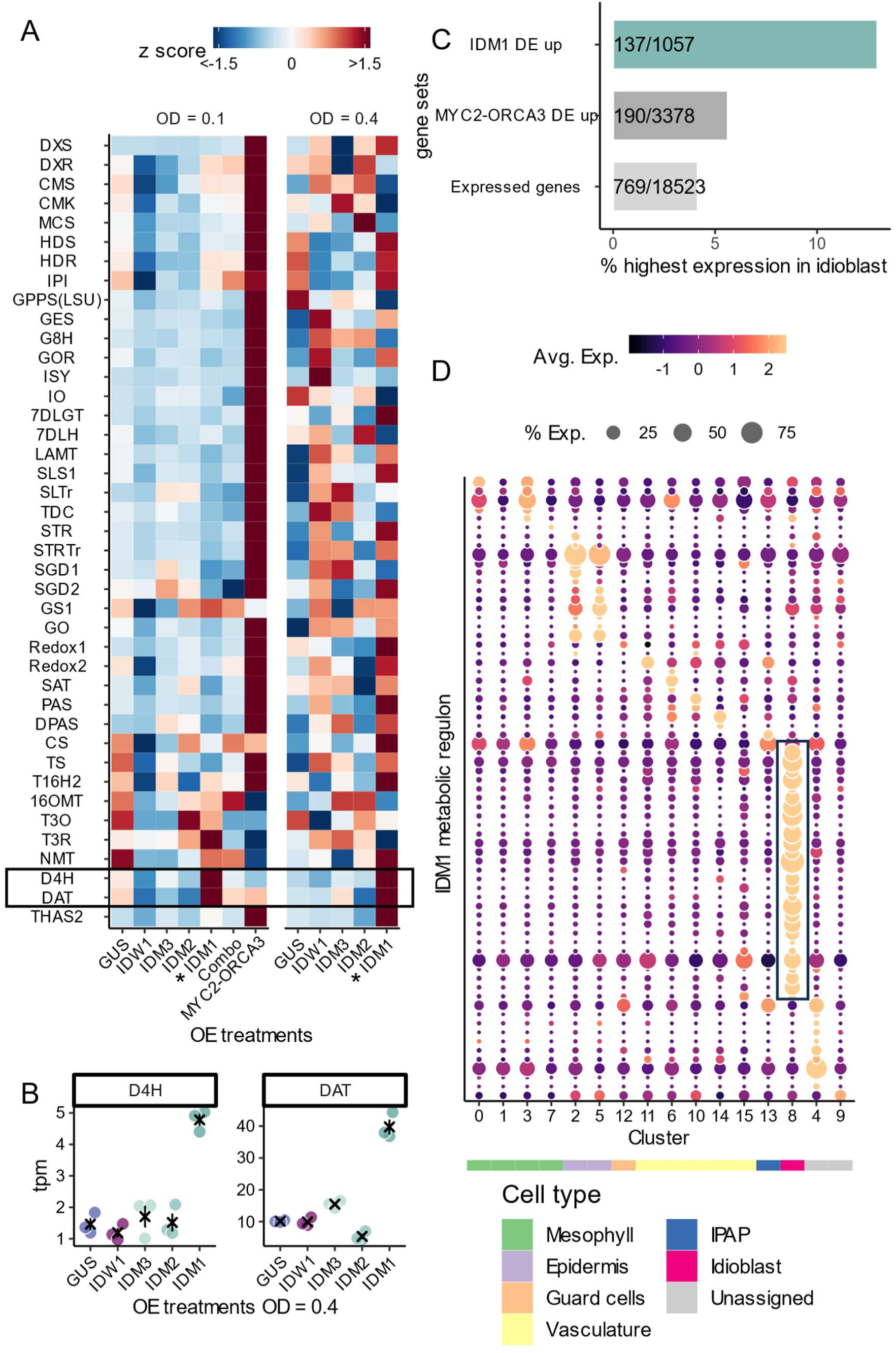
Idioblast MYB1 (IDM1) activates the expression of D4H and DAT, as well as an idioblast-specific transcriptional program. A. Gene expression heatmap of the MIA biosynthetic genes across overexpression treatments. Each row is a biosynthetic gene or transporter, ordered from upstream to downstream. Color scale represents scaled expression (z score). Combo: the combinatory treatment in which IDW1 and IDM1/2/3 are co-infiltrated. B. Mean separation plots showing expression levels of D4H and DAT (in units of transcripts per million) in the 0.4 OD treatments. Each data point is a biological replicate. Error bars represent average and standard error. Black × indicates average. C. Bar graph showing percentage of genes that are most highly expressed in the idioblast. Expressed genes: all 18,523 expressed genes in this single cell multiome dataset. MYC2-ORCA3: 3,378 differentially expressed genes that are upregulated in the MYC2-ORCA3 overexpression treatment. IDM1: 1,057 differentially expressed genes that are upregulated in the 0.4 OD overexpression IDM1 treatment. D. Gene expression heatmap of IDM1 metabolic regulon (see also Supplementary Table 11). Color scale shows the average scaled expression of each gene at each cell cluster. Dot size indicates the percentage of cells where a given gene is detected. The predicted cell type for each cell cluster is annotated by the color strip below the x-axis. Box highlights genes specifically expressed in the idioblast.

We found that IDM1 continued to activate both D4H and DAT at 0.4 OD (Fig. 5A, B), resulting in even higher fold changes (3.2-fold and 3.9-fold increase relative to GUS control of the corresponding experiment, respectively).

In addition to D4H and DAT, we found that genes differentially upregulated by IDM1 were enriched for idioblast expression (Fig. 5C). Among all 18,523 expressed genes in the single cell multiome dataset, only 769 (4% of 18,523) were most highly expressed in the idioblast. Similarly, among 3,378 differentially upregulated genes in the MYC2-ORCA3 treatment, 190 (5.6% of 18,523) were most highly expressed in the idioblast. In contrast, among 1,057 differentially upregulated genes in the IDM1 treatment at 0.4 OD, 137 (13% of 1,057) were most highly expressed in the idioblast, representing a 3.3-fold enrichment over the background of all expressed genes (*p* < 2.2 × 10^-16^, χ-squared test). IDM1 also activated IDW1 and IDM2/3, the three other idioblast specific WRKY and MYB TF candidates described above (Supplementary Fig. 10C). Gene set enrichment analyses revealed that, similar to MYC2-ORCA3, IDM1 upregulated genes were enriched for gene families relevant to specialized metabolism (transporters, cytochrome P450s, alcohol dehydrogenases, and 2-OG-dependent oxygenases). For example, among all expressed genes, 0.27% of them are annotated as alcohol dehydrogenases, whereas 0.65% and 1.3% of MYC2-ORCA3 and IDM1 upregulated genes were annotated as alcohol dehydrogenase, respectively. These IDM1 upregulated genes that are potentially relevant to specialized metabolism were designated as the IDM1 metabolic regulon (*n* = 61 genes, Supplementary Table 11). We found that 44% (27/61) of the IDM1 metabolic regulon were specifically expressed in the idioblast (Fig. 5D), more than 10-fold enrichment over the background of all expressed genes (background = 4% of 18,523 expressed genes, *p* < 2.2 × 10^-16^, χ-squared test). Taken together, these observations suggest IDM1 regulates an idioblast specific transcriptional program, which includes the MIA biosynthetic genes D4H and DAT, as well as other gene families potentially involved in natural product biosynthesis.

We next tested whether IDM1 could directly activate the expression of D4H and DAT using reporter transactivation assays (Supplementary Fig. S11-12). We first confirmed that IDM1 is localized to the nucleus (Supplementary Fig. S11A-D). To construct reporters, we fused accessible chromatin regions upstream of DAT (Fig. 3B) and D4H (Supplementary Fig. 11E) to a minimal 35S promoter driving an DsRed reporter. On the same plasmid, a GFP internal control was included, which is driven by the constitutive Arabidopsis UBQ1 promoter (Supplementary Fig. 11F). We then performed the transactivation assay by co-infiltrating an agrobacterium strain carrying 35S:IDM1 (Supplementary Fig. 11G) and a strain carrying the reporter construct for either DAT or D4H. We observed conspicuous DsRed^+^ cells in infiltrated petals for both DAT and D4H reporters (Supplementary Fig. 12A, B, E, F). In contrast, no DsRed^+^ cells in petals could be observed when either reporter was infiltrated alone (Supplementary Fig. 12C, D, G, H,). These observations were confirmed by pixel intensity quantifications using ImageJ ^43^. High red to green pixel intensity ratio was only detected when 35S:IDM1 and one of DAT or D4H reporters were co-infiltrated (Supplementary Fig. 12I). In contrast, low red to green ratio was detected when the reporter was infiltrated without 35S:IDM1. IMD1 could not transactivate a reporter construct that did not contain the DAT or D4H accessible chromatin regions (Supplementary Fig. 12J-O), which was confirmed by pixel intensity quantifications (Supplementary Fig. 12I). Taken together, these results strongly suggest that IDM1 is a direct activator of D4H and DAT, and the idioblast specific expression of IDM1 contributes to the idioblast specific expression of D4H and DAT.

## Discussion

The cell-type-specific expression patterns of MIA biosynthetic genes in *C. roseus* are well documented ^5,7,8^. In this study, using single cell multi-omics datasets, we discovered the first reported idioblast specific TF (CrIDM1) that regulates late-stage vinblastine biosynthetic genes (D4H and DAT). Although several TFs that regulate MIA biosynthesis have been characterized ^11–20,28,29^, how the exquisite cell-type specific regulation is achieved for this pathway remains unclear. We generated the first single cell multiome dataset for *C. roseus* leaves to investigate gene regulation of the MIA pathway at single cell resolution. Not only did we recapitulate the cell-type specific expression pattern of the pathway, but we also catalogued a dictionary of *cis*-regulatory elements associated with MIA biosynthetic genes. We showed that among previously studied TFs pertinent to the MIA pathway, only BIS1/2/3 and ORCA4 were co-expressed with their target genes at the cell type level (Fig. 2D, Supplementary Fig. 7A), suggesting BIS1/2/3 and ORCA4 contribute to the cell-type specific expression pattern of the MIA biosynthetic pathway.

There is little information on how the pathway is regulated beyond Catharanthine Synthase (Fig. 2D, Fig. 5A). The late-stage MIA biosynthetic genes were marked with cell-type specific ATAC-seq peaks, suggestive of coordinated regulation at the chromatin level (Fig. 3A). Epidermis marker peaks (Fig. 3B) were enriched for homeodomain, ERF, and MYB binding motifs (Supplementary Fig. 7B). Members of the above-mentioned TF families have been reported to regulate other specialized metabolism pathways, such as anthocyanin ^30^, cuticle ^44^, suberin ^45^, and glucosinolate ^46^ in other species. We speculate that yet unidentified homeodomain, ERF, and MYB TFs may contribute to the cell type specific expression of MIA biosynthetic genes in epidermis. The dataset generated in this study can be used to mine and characterize additional metabolic regulators that operate specifically in the epidermis.

We found that WRKY and MYB motifs were overrepresented among idioblast marker peaks (Fig. 3C). Paired with gene co-expression analyses across cell clusters, we narrowed down our candidates to a single WRKY (IDW1) and three MYB TFs (IDM1/2/3) (Fig. 4). While candidate TFs can be identified from gene expression data alone ^8,35^, we demonstrated that cell-type specific chromatin accessibility profiles allowed us to identify putative cell-type specific *cis*-regulatory elements and the corresponding TF families using motif enrichment (Fig. 4C, Supplementary Fig. 7A), which in turn pin-pointed TF candidates that most likely activate target genes in a cell-type specific manner.

Overexpression and reporter transactivation assays demonstrated that IDM1 is a novel idioblast specific regulator for D4H and DAT (Fig. 5). IDM1 binds the accessible chromatin regions upstream of D4H and DAT and activates their expression (Supplementary Fig. 10-11). Recently, a GATA family TF, GATA1 was reported to activate the expression of late vinblastine biosynthetic genes in de-etiolating seedlings, including T16H2, T3O, T3R, D4H and DAT ^28^. However, we found that GATA1 was only expressed in the mesophyll of the leaf in our single cell dataset (Supplementary Fig. 7A), suggesting GATA1 is likely not responsible for the cell-type specific patterns of the late-stage pathway. In contrast, IDM1 is expressed exclusively in the idioblast, and thus it contributes to the idioblast specific expression of D4H and DAT. Since IDM1 is also JA-inducible (Supplementary Fig. 7C), IDM1 may also mediate JA-dependent activation of D4H and DAT.

In addition to D4H and DAT, IDM1 activates an idioblast metabolic regulon (Fig. 5C, D). Gene sets such as transporters, cytochrome P450, alcohol dehydrogenase, and 2-OG dependent oxygenase are strongly enriched in IDM1 upregulated genes, suggesting that IDM1 is a *bona fide* metabolic regulator. The IDM1 metabolic regulon is highly enriched for idioblast specific expression (Fig. 5D), suggesting other targets of IDM1 may play a role in the biosynthesis of vinblastine or other alkaloids in the idioblast. IDM1 activates IDW1 and IDM2/3, which did not appear to activate the MIA pathway, at least in the experimental conditions we tested (Fig. 5A). IDW1 and IDM2/3 might regulate other biological processes in the idioblast, which may be important for the specialization of these rare cells. Even after decades of focused research, the final steps of the *C. roseus* MIA biosynthetic pathway remains an enigma. The discovery of IDM1 as a regulator of the late stages of MIA biosynthesis and access to an idioblast-specific gene regulatory network will expedite completion of this 40-plus step biosynthetic pathway with important human-health implications.

## Methods

### Nuclei isolation and single cell library preparation

*Catharanthus roseus* (cultivar “Sunstorm Apricot”) plants were grown in under a 14-hr photoperiod at 22 °C. Mature, fully expanded leaves were sampled from 8-10-week-old plants. Nuclei isolation was performed as described previously ^47^ with 0.01% Triton-X-100 in the nuclei isolation buffer. Around 0.3-0.5 g of leaves were chopped vigorously on ice on a petri dish in nuclei isolation buffer for exactly 2 min. The lysate was filtered through 100 µm and 40 µm sieves, before passing through a 20 µm strainer twice. Nuclei were stained with 4’,6-diamidino-2-phenylindole (DAPI) and sorted using a Moflo Astrios EQ flow cytometer at the UGA Cytometry Shared Resource Laboratory. At least 100,000 nuclei were sorted into 500 µL of nuclei buffer (part of 10x Genomics Single Cell Multiome Kit). Sorted nuclei were pelleted by centrifugation at 200 g for 5 min and resuspended in 50 µL nuclei buffer. The integrity of the nuclei was visually inspected using a fluorescence microscope (Supplementary Fig. 1B-I). Multiome libraries were constructed using the 10x Genomics Single Cell Multiome Kit, according to manufacturer’s instruction.

### Single nuclei RNA-seq processing

Single nuclei RNA-seq libraries were processed using Cutadapt (v3.5) ^48^ with the following parameters: -q 30 -m 30 --trim-n -n 2 -g AAGCAGTGGTATCAACGCAGAGTACATGGG -a “A{20}”. The pairing of the reads was restored using SeqKit (v0.16.1) *pair* ^49^. Paired reads were aligned and quantified using STARsolo ^50^, with the following parameters: --runThreadN 24 --alignIntronMax 5000 --soloUMIlen 12 --soloCellFilter EmptyDrops_CR --soloFeatures GeneFull --soloMultiMappers EM --soloType CB_UMI_Simple, and --soloCBwhitelist using the latest 10x Genomics whitelist of multiome barcodes. Gene-barcode matrices were analyzed with Seurat (v4) ^51^ for downstream analysis. Removal of low-quality nuclei and suspected multiplets was performed using the distributions of UMI counts and detected genes (Supplementary Fig. 2).

### Single nuclei RNA-seq analyses

Biological replicates were integrated using the ‘IntegrateData()’ function in Seurat using the top 3,000 variable genes. Uniform manifold approximation and projection (UMAP) were performed after a principal component analysis (PCA) using the following parameters: dims = 1:30, min.dist = 0.001, repulsion.strength = 1, n.neighbors = 15, spread = 5. Clustering of cells was performed with a resolution of 0.5. For cell type classification, we used a manually curated marker gene list for mesophyll, epidermis, guard cells, and vasculature (Supplementary Table 5), using previously established marker genes from Arabidopsis ^52,53^ and *C. roseus* ^5–8^. For dot-plot style expression heat maps, average expression of genes was calculated as the average Z-score of log-transformed normalized expression values across cell clusters and cell types. Dot sizes indicated the percentage of cells where a given gene is expressed (> 0 reads) in each cell type or cell cluster.

### Single nuclei ATAC-seq processing

Single nuclei ATAC-seq data were processed using the 10x Genomics Cell Ranger ARC pipeline (https://www.10xgenomics.com/software). For initial quality control and nuclei filtering, the ‘atac_peaks.bed’ files from the Cell Ranger ARC output were used. The peak bed files for the three biological replicates were sorted and merged using BEDTools (v2.30) *merge* ^54^. This common set of peaks was used to process all three biological replicates. The ‘atac_fragments.tsv.gz’ files from the Cell Ranger ARC output were used for downstream analyses using Signac (v1.6.0) ^55^ and Seurat (v4) ^51^. Nuclei were filtered for > 1000 peaks/nuclei, > 2000 fragments/nuclei, and fraction of fragments in peaks > 0.25. For data integration, the replicates were merged first, then integrated using the ‘IntegrateEmbeddings()’ function in Signac using the “lsi” dimension reduction. Integration with the gene expression assay was performed by first filtering for shared nuclei in both gene expression and chromatin assays, after which the integrated ATAC-seq object was adjoined to the integrated RNA-seq object as a chromatin assay. By doing so, the cell cluster and cell type assignment information is transferred to the ATAC-seq assay. Fragment files were split into separate files for each cell cluster and converted to bed files. Peak calling at each cell cluster performed using MACS2 (v2.2.7.1) ^23^ using the following parameters: -f BED -g 444800000 (80% of the genome assembly size was set as the effective mappable genome size) --nomodel --broad. The resultant peak files were sorted and merged to be used as the features in the chromatin accessibility assay. These peaks were used as “ATAC-seq peaks” in all downstream analyses. UMAP visualization (Fig. 1C) for ATAC-seq was performed using the following parameters: reduction = “lsi”, dims = 2:30, min.dist = 0.001, repulsion.strength = 1, n.neighbors = 30, spread = 1. Joint UMAP visualization was done using the ‘FindMultiModalNeighbors()’ functions in Signac. ATAC-seq coverage around genes (Supplementary Fig. 4) and peaks (Supplementary Fig. 5) was calculated and visualized using deepTools (v3.5.1) ^56^.

### DAP-seq library construction and processing

The coding sequence of *ORCA3* and *ORCA4* were synthesized and cloned into pIX-Halo ^24^, downstream and in frame with the halo tag. *In vitro* gene expression was performed using Promega TnT SP6 High-Yield Wheat Germ Protein Expression System. Each *in vitro* gene expression reaction was spiked with 200 ng of a pIX-RFP plasmid, such that the gene expression reaction can be monitored using RFP fluorescence. Genomic data libraries were constructed from genomic DNA isolated from mature leaves of 8-10-week-old *C. roseus* plants using a KAPA HyperPrep Kit, after the genomic DNA was sheared to 200-bp with a Covaris ultrasonicator at the UGA Genomics and Bioinformatics Core. The full volume of gene expression reaction was combined with 40 ng of gDNA library and 10 µL of Promega Halo-beads for each affinity reaction. Bead-bound DNA was recovered by heating the affinity reaction to 95°C for 5 min. Indexing PCR was performed with 13 cycles, and the libraries were sequenced in paired-end 50-bp format (Supplementary Table 2).

Sequencing adapters were trimmed with Cutadapt (v3.5) ^48^, after which reads were aligned to the *C. roseus* v3 genome ^8^ using BWA mem (v0.1.17) ^57^. Peak calling was performed with MACS2 (v2.2.7.1) using the following parameters: -g 444800000 (80% of the genome assembly size was set as the effective mappable genome size), using the bam file of the halo tag control as the background file. DAP-seq coverage around peaks (Supplementary Fig. 6E, F) was calculated and visualized using deepTools (v3.5.1). Putative target genes were assigned using BEDTools (v.2.30) *closest*, with the -d parameter selected. Genes overlapping or within 2-kb of a DAP-seq peak were designated as a putative target gene. Accessible DAP-seq peaks were defined as DAP-seq peaks overlapping or within 100-bp to either ends of an ATAC-seq peak (Supplementary Fig. 6D). DNA sequence of DAP-seq peaks were extracted using BEDTools (v.2.30) *getfasta* and subjected to *de novo* motif discovery using MEME (v5.4.1) ^25^: using the following parameters: -dna -revcomp -mod anr -nmotifs 10 -minw 5 -maxw 12 -evt 0.01.

### Marker peak and motif overrepresentation analyses

Marker peaks for epidermis and idioblast were detected using the ‘FindMarkers()’ function in Seurat after setting the default assay of the multiome object to chromatin accessibility, using the following parameters: only.pos = T, test.use = “LR”, min.pct = 0.05, latent.vars = ‘nCount_peaks’, group.by = “cell_type”. Only peaks with adjusted p values < 0.05 were used for downstream analyses. For motif enrichment analysis, position weight matrices were obtained using the ‘getMatrixSet()’ function in Signac, using the following parameters: x = JASPAR2020 ^27^, opts = list(collection = “CORE”, tax_group = ‘plants’, all_versions = FALSE). These motifs were added to the multiome object using the ‘AddMotifs()’ function in Signac. Overrepresented motifs were identified using the ‘FindMotifs()’ function in Signac.

### Gene co-expression analyses

Gene co-expression analysis by graph-based clustering was performed as previously described ^33^. The top 3,000 most variable genes were used for gene-wise correlation. Pairwise Pearson correlation was performed to generate an edge table, which was filtered for *r* > 0.75. Graph-based clustering was performed with a resolution parameter of 4.

### Overexpression assays

Coding sequences of *IDW1* and *IDM1/2/3* were cloned in between the 35S promoter and 35S terminator and transformed into *Agrobacterium tumefaciens* strain GV3101. We used previously published MYC2 and ORCA3 overexpression constructs ^14^. Transient expression experiments were done on *C. roseus* (cultivar “Little Bright Eyes”) petals. Infiltration was done as previously described ^58^. Two days before the infiltration, all open flowers were removed. Two sets of experiments were performed. In the first set, individual strains were infiltrated at 0.1 OD and the total OD was adjusted to 0.4 using the control agrobacterium strain carrying GUS. In the second set, all strains were infiltrated at 0.4 OD. Two days after the infiltration, infiltrated petals were harvested and stored in a −80 freezer until RNA extraction.

### RNA-seq analysis for overexpression samples

Sequencing adapters were trimmed from petal RNA-seq libraries using Cutadapt (v3.5) ^48^. Adapter trimmed libraries were pseudo-aligned and quantified using kallisto (v0.48) ^59^, with the --plaintext option turned on. When the appropriate strandedness parameter was used, pseudo-alignment rate ranged from 86.2% to 89%. Differential gene expression analyses were performed using DESeq2 (v.1.34.0) ^60^, using the GUS treatment with of the corresponding experiment as control. Genes with adjusted p values < 0.05 were taken as differentially expressed genes.

### Reporter transactivation assays

The reporter transactivation assays were performed in a two-component format: a reporter component and an overexpression component. Genetic parts used in reporter assays were amplified from a vector tool kit for plant molecular biology ^61^. The accessible chromatin regions immediately upstream of D4H (Supplementary Fig. 11A) and DAT (Fig. 3B) were cloned upstream of a 35S minimal promoter (Supplementary Fig. 11B), which controls the expression of DsRed reporter. On the same plasmid, a GFP internal control driven by the Arabidopsis UBQ1 promoter was also included. The overexpression component was an agrobacterium GV3101 strain carrying 35S:IDM1 (Supplementary Fig. 11C), the same construct used in overexpression assays. As in the transient expression assays described above, experiments were done on *C. roseus* (cultivar “Little Bright Eyes”). Two days after the infiltration, petals were imaged using a fluorescent microscope. Pixel intensity was quantified using ImageJ ^43^.

## Data Availability

All sequencing data associated with this study are available at the National Center for Biotechnology Institute Sequence Read Archive BioProject PRJNA1098712. Seurat objects for single nuclei multiome experiment and gene expression matrices are available via the online digital repository figshare (to be made public upon publication). Plasmid maps are available at Zenodo (https://zenodo.org/records/11036874).

## Code Availability

All custom codes used to generate figures can be found at https://github.com/cxli233/Catharanthus_multiome.

## Acknowledgements

This project was supported by the Georgia Research Alliance (C.R.B.) and Georgia Seed Development (C.R.B.), National Science Foundation MCB-2309665 (C.R.B and C.L.), and the Max Planck Gesellschaft (S.E.O and L.C.). Sequencing was performed at Biomarker Technologies (BMK) GmbH, Max-Planck Institute for Biochemistry Next Generation Sequencing Facility, Novogene, Texas A&M AgriLife Research: Genomics and Bioinformatics Service, and University of Georgia Genomics and Bioinformatics Core (GGBC, UG Athens, GA, RRID:SCR_010994). The author would like to acknowledge Julie Nelson at University of Georgia Cytometry Shared Resource Laboratory for assistance with flow cytometry, Dr. Alexandre P. Marand for suggestions on nuclei isolation protocol development, Dr. Robert J. Schmitz for advice on ATAC-seq analyses, and Dr. Alain Goossens for providing plasmids.

## Author contributions

C.R.B., S.E.O., and C.L. designed the study. C.L. generated single cell multiome and DAP-seq datasets. J.C.W. assisted with single cell library preparation and quality control. C.L. and S.L.J performed molecular cloning and transactivation assays. M.C. performed molecular cloning and overexpression assays, and together with L.C. generated overexpression samples and RNA-seq datasets. C.L., J.C.W, B.V., and J.P.H performed data analyses. C.L. wrote the manuscript with input from all authors.

