## Supplementary Figure and Legends for "Cell-type aware regulatory landscapes governing monoterpene indole alkaloid biosynthesis in the medicinal plant *Catharanthus roseus*"

**Supplementary Figures**

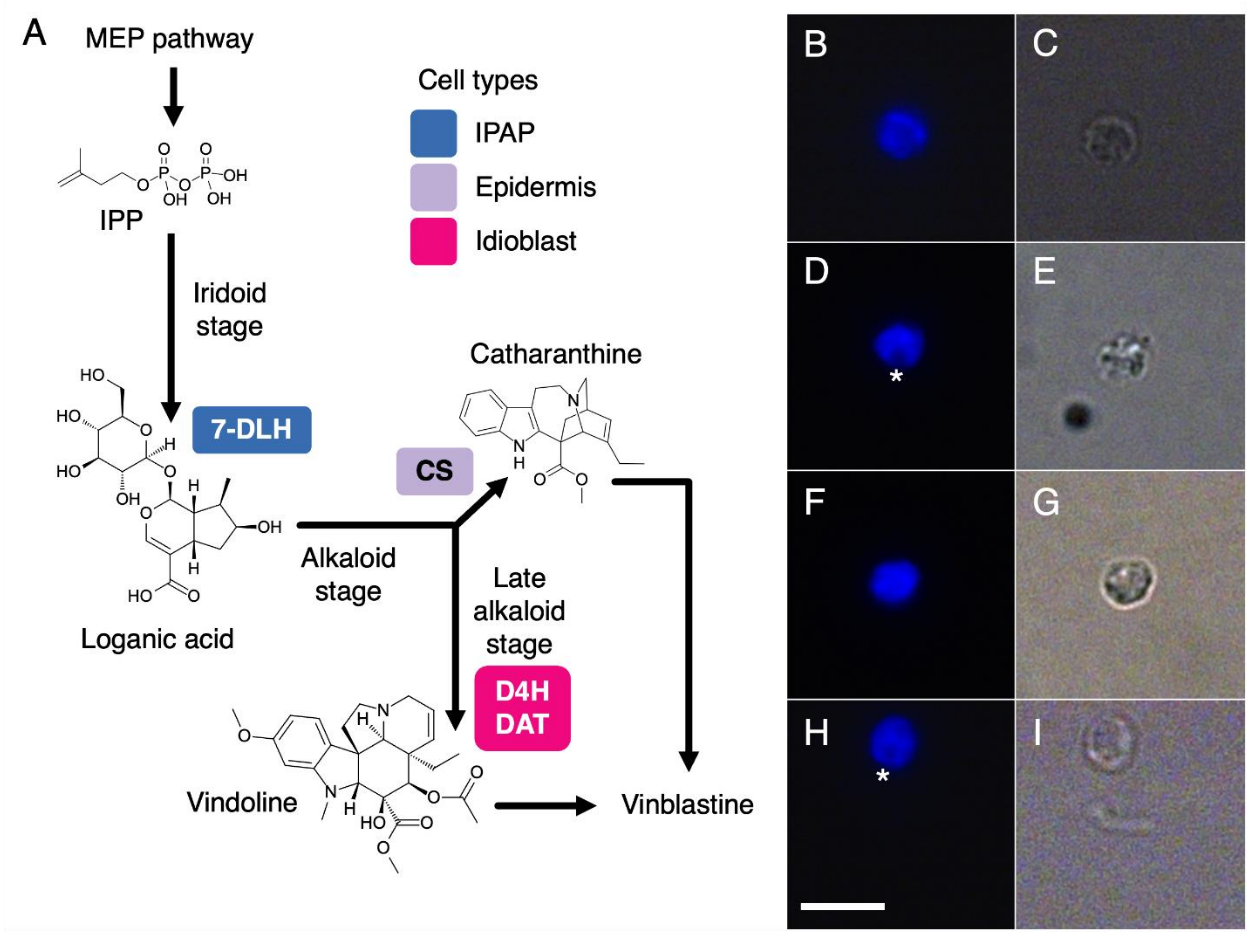


**Supplementary Fig. 1. Schematics of the pathway and isolated nuclei.**

A. Simplified schematics of the MIA pathway in *C. roseus*. Structures and names of key intermediates are shown. Key enzymatic steps are shown and highlighted by the cell types they are expressed in. IPAP: internal phloem associated parenchyma. See also Supplementary Table 1 for gene names.

B-I. Intact, isolated nuclei from *C. roseus* leaves used for single cell multiome experiments.
B, D, F, H. DAPI fluorescence.
C, E, G, I. Bright field.
Asterisks: nucleolus. Bar: 10 μm.

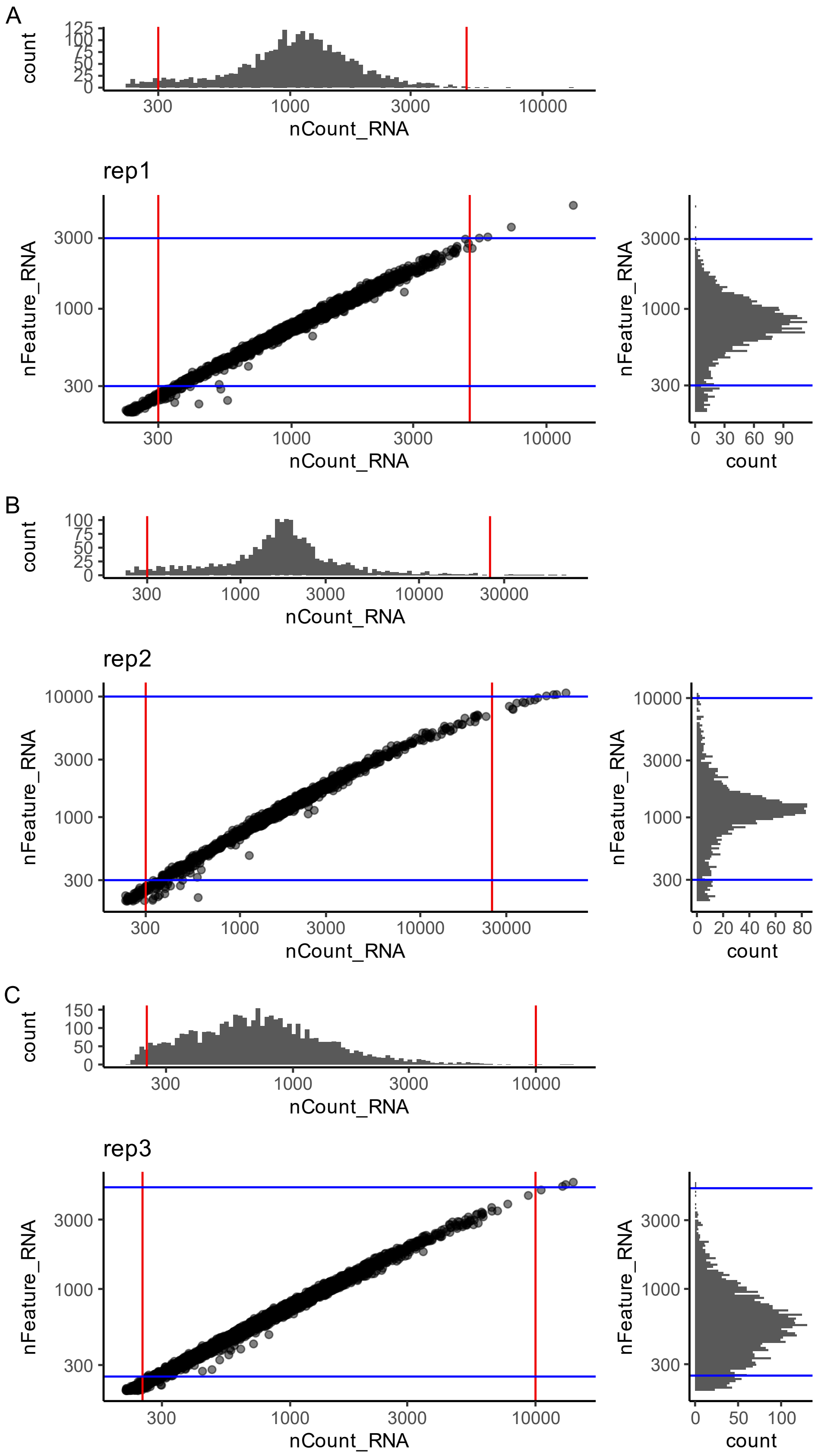
A
count
125
100
75
50
25
0
300
1000
3000
10000
nCount_RNA
rep1
nFeature_RNA
3000
1000
300
nCount_RNA
count
0 30 60 90
B
100
75
50
25
0
30000
rep2
10000
0 20 40 60 80
C
150
rep3
0 50 100

**Supplementary Fig. 2. Evaluation and quality control for the gene expression assay of single cell multiome dataset.**

A-C: scatter plots of nCount_RNA (UMI count) and nFeature_RNA (number of detected genes) for three biological replicates. Both axes are in $log_{10}$ scale. The distribution of UMI count and gene count are shown above and to the right of the scatter plots, respectively. Only nuclei between the red and blue lines are used for downstream analyses. See also Supplementary Table 3.

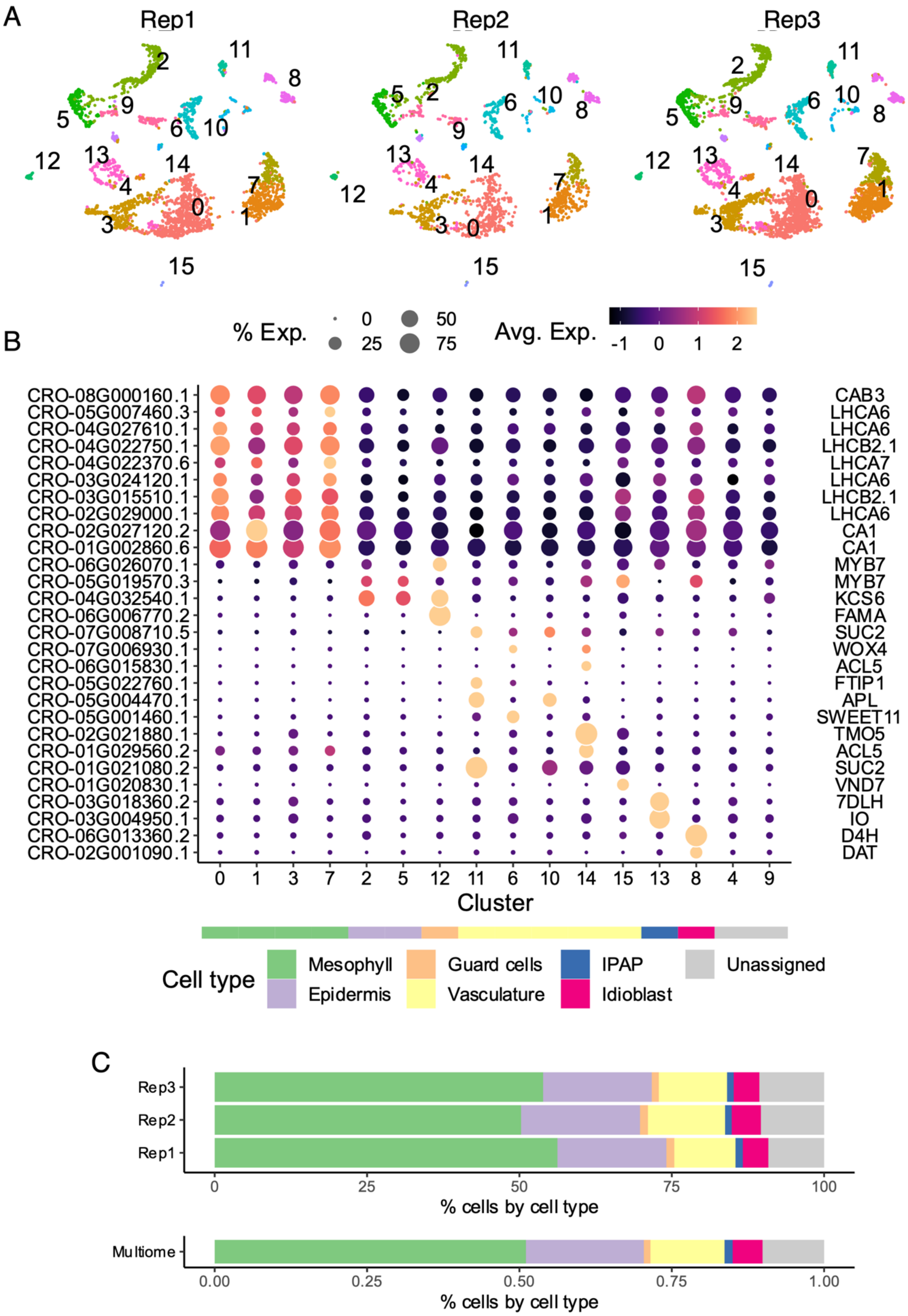
A
Rep1
Rep2
Rep3
B
% Exp.
0
25
50
75
Avg. Exp.
-1
0
1
2
CRO-08G000160.1
CRO-05G007460.3
CRO-04G027610.1
CRO-04G022750.1
CRO-04G022370.6
CRO-03G024120.1
CRO-03G015510.1
CRO-02G029000.1
CRO-02G027120.2
CRO-01G002860.6
CRO-06G026070.1
CRO-05G019570.3
CRO-04G032540.1
CRO-06G006770.2
CRO-07G008710.5
CRO-07G006930.1
CRO-06G015830.1
CRO-05G022760.1
CRO-05G004470.1
CRO-05G001460.1
CRO-02G021880.1
CRO-01G029560.2
CRO-01G021080.2
CRO-01G020830.1
CRO-03G018360.2
CRO-03G004950.1
CRO-06G013360.2
CRO-02G001090.1
CAB3
LHCA6
LHCA6
LHCB2.1
LHCA7
LHCA6
LHCB2.1
LHCA6
CA1
CA1
MYB7
MYB7
KCS6
FAMA
SUC2
WOX4
ACL5
FTIP1
APL
SWEET11
TMO5
ACL5
SUC2
VND7
7DLH
IO
D4H
DAT
0 1 3 7 2 5 12 11 6 10 14 15 13 8 4 9
Cluster
Cell type
Mesophyll
Epidermis
Guard cells
Vasculature
IPAP
Idioblast
Unassigned
C
Rep3
Rep2
Rep1
0 25 50 75 100
% cells by cell type
Multiome
0.00 0.25 0.50 0.75 1.00
% cells by cell type

**Supplementary Fig. 3. Cell clustering and cell type marker gene analyses.**

A. UMAP, as shown in Fig. 1B, split by biological replicates.

B. Gene expression heatmap of marker genes across cell clusters. Color scale shows the average scaled expression of each gene at each cell cluster. Dot size indicates the percentage of cells where a given gene is detected. The predicted cell type for each cell cluster is annotated by the color strip below the x-axis. Arabidopsis and *C. roseus* gene symbols shown on the right (see also Supplementary Table 4).

C. A stacked bar plot showing percentage of cells in each type. The color code is the same as (B).

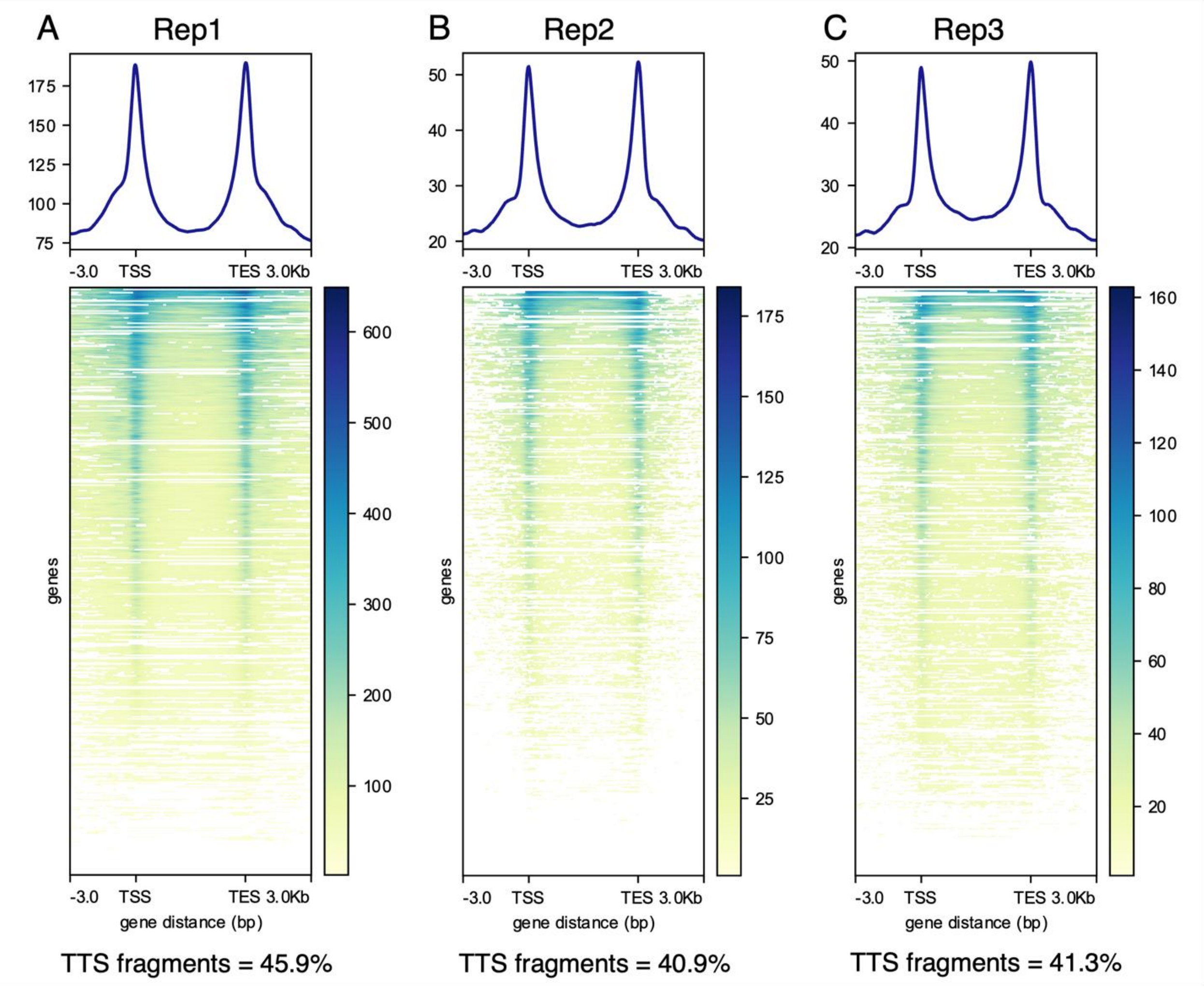


**Supplementary Fig. 4. Evaluation of chromatin accessibility profiles around genes.**

A-C. Metagene plot (top) and heatmap (bottom) showing coverage of ATAC-seq fragments around genes for each biological replicate. In the heatmap, each row is a gene. TSS. Transcription start site. TES. Transcription end site. The percentage of fragments overlapping TSSs are listed below the heatmap. See also Supplementary Table 5.

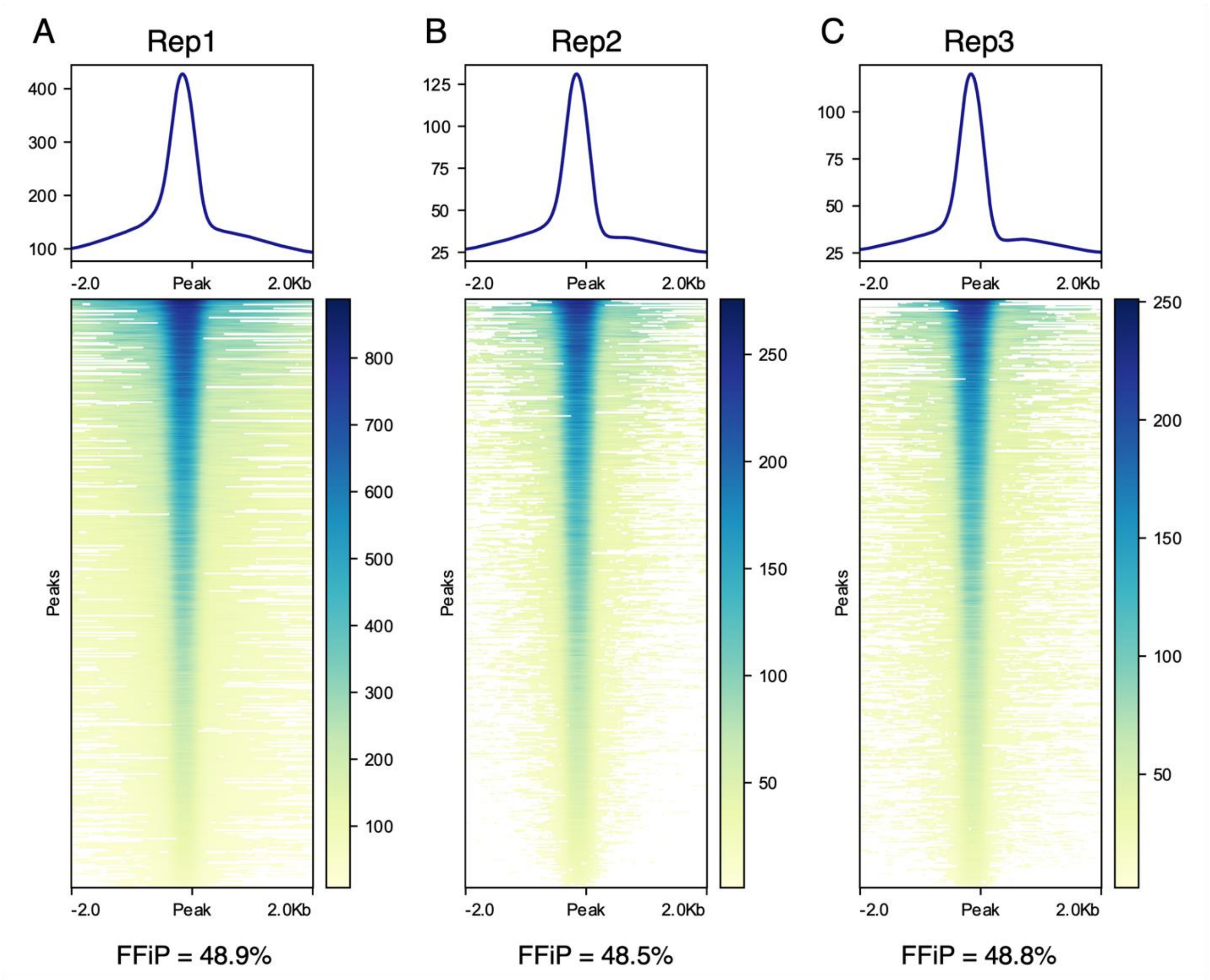


**Supplementary Fig. 5. Evaluation of chromatin accessibility profiles around ATAC-seq peaks.**

A-C. Metagene plot (top) and heatmap (bottom) showing coverage of ATAC-seq fragments around ATAC-seq peaks for each biological replicate. In the heatmap, each row is an ATAC-seq peak. The percentage of fragments overlapping peaks are listed below the heatmap. FFiP: fraction of fragments in peaks. See also Supplementary Table 5.

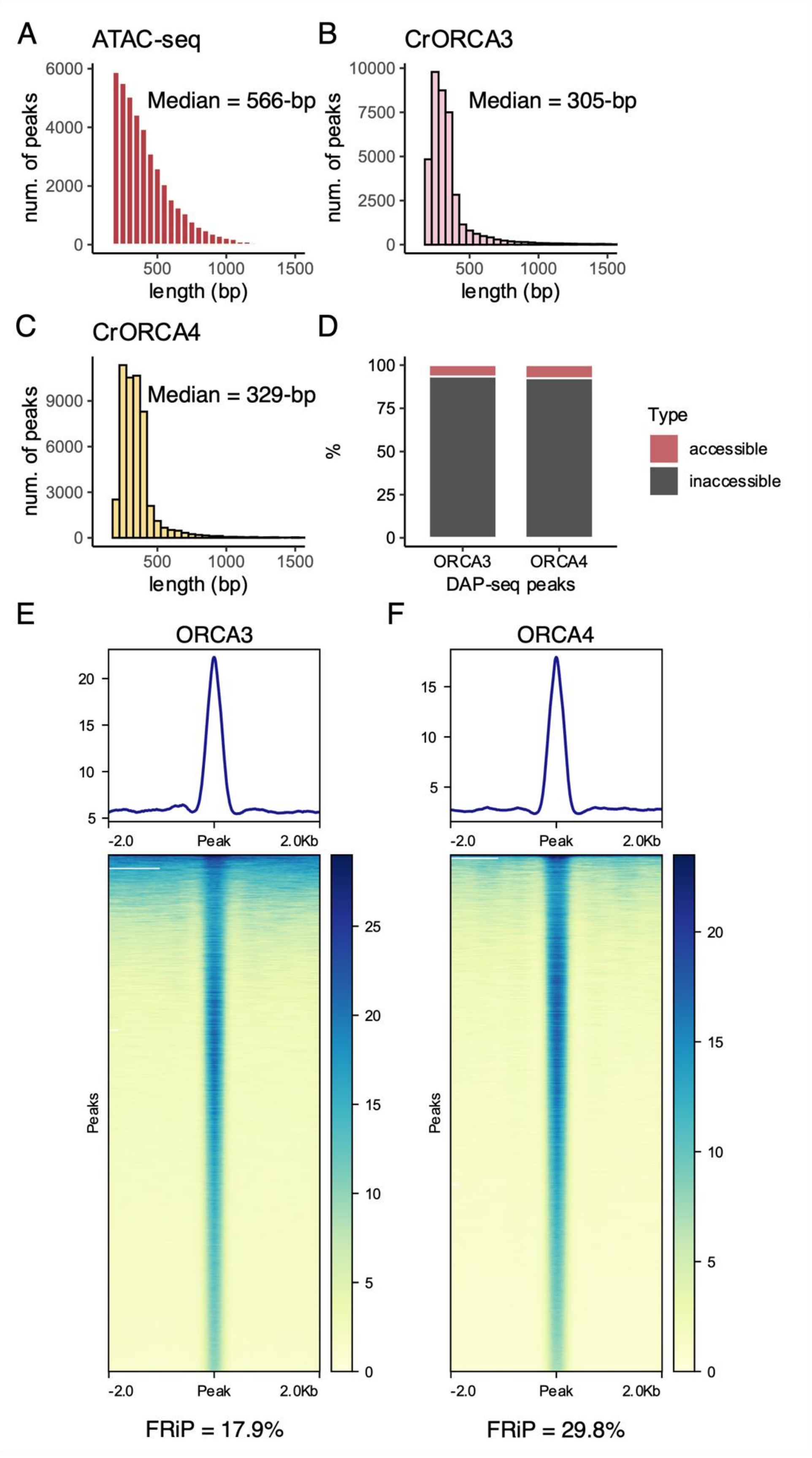

A
ATAC-seq
Median = 566-bp
num. of peaks
length (bp)
B
CrORCA3
Median = 305-bp
num. of peaks
length (bp)
C
CrORCA4
Median = 329-bp
num. of peaks
length (bp)
D
%
Type
accessible
inaccessible
ORCA3
ORCA4
DAP-seq peaks
E
ORCA3
-2.0
Peak
2.0Kb
Peaks
FRiP = 17.9%
F
ORCA4
-2.0
Peak
2.0Kb
Peaks
FRiP = 29.8%

**Supplementary Fig. 6. Evaluation of DAP-seq peaks.**

A-C. Histograms showing distribution of peaks lengths. A. ATAC-seq peaks; B. ORCA3 DAP-seq peaks. C. ORCA4 DAP-seq peaks.

D. Stacked bar plots showing percentages of DAP-seq peaks that are accessible (see Methods).

E-F. Metagene plot (top) and heatmap (bottom) showing coverage of DAP-seq reads around DAP-seq peaks for each ORCA3 (E) and ORCA4 (F). In the heatmap, each row is a DAP-seq peak. The percentage of reads overlapping peaks are listed below the heatmap. FRiP: fraction of reads in peaks. See also Supplementary Table 6.

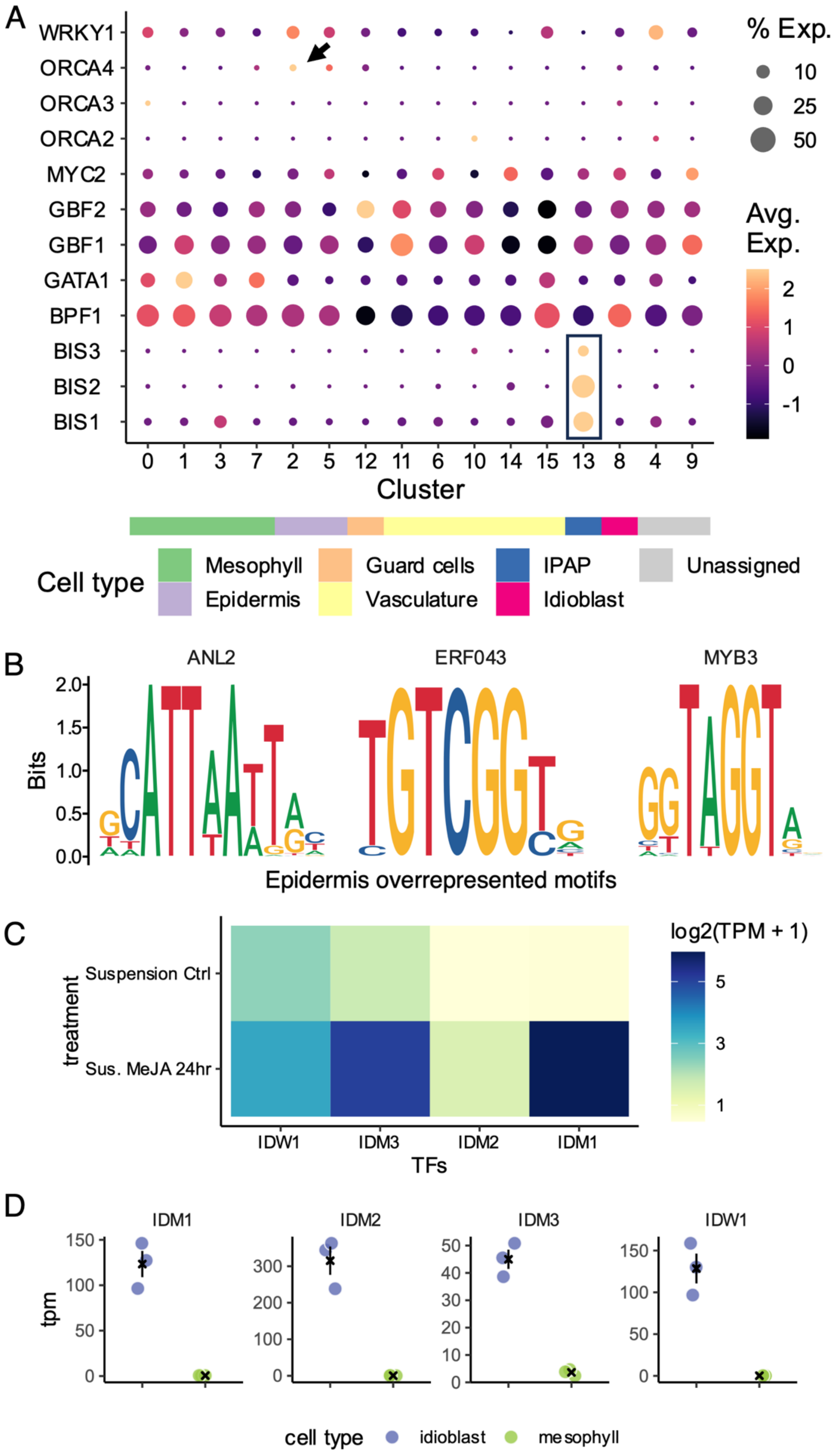
A
WRKY1
ORCA4
ORCA3
ORCA2
MYC2
GBF2
GBF1
GATA1
BPF1
BIS3
BIS2
BIS1
0 1 3 7 2 5 12 11 6 10 14 15 13 8 4 9
Cluster
% Exp.
10
25
50
Avg. Exp.
2
1
0
-1
Cell type
Mesophyll
Epidermis
Guard cells
Vasculature
IPAP
Idioblast
Unassigned
B
ANL2
ERF043
MYB3
Bits
2.0
1.5
1.0
0.5
0.0
Epidermis overrepresented motifs
C
treatment
Suspension Ctrl
Sus. MeJA 24hr
IDW1
IDM3
IDM2
IDM1
TFs
log2(TPM + 1)
5
3
1
D
IDM1
IDM2
IDM3
IDW1
tpm
cell type
idioblast
mesophyll

**Supplementary Fig. 7. Gene expression and motif enrichment analyses regarding known TFs, epidermis marker peaks, and idioblats TF candidates.**

A. Gene expression heatmap of previously studied TFs across cell clusters. Color scale shows the average scaled expression of each gene at each cell cluster. Dot size indicates the percentage of cells where a given gene is detected. The predicted cell type for each cell cluster is annotated by the color strip below the x-axis. Arrow highlights the epidermis specific expression of ORCA4. Box highlights the IPAP-specific expression of BIS1/2/3. See also Supplementary Table 7.

B. TF binding motifs overrepresented among epidermis marker peaks.

C. $\log_2$ transformed transcripts per million (TPM) values for idioblast TF candidates (see also Fig. 4C) in a methyl-jasmonate treatment experiment. Ctrl. Control. Sus. MeJA 24 hr. Suspension culture treated with methyl-jasmonate for 24 hrs.

D. Transcripts per million values for idioblast TF candidates in a flow cytometry sorted protoplast experiment. Error bars indicate average and standard error. Black × indicates average.

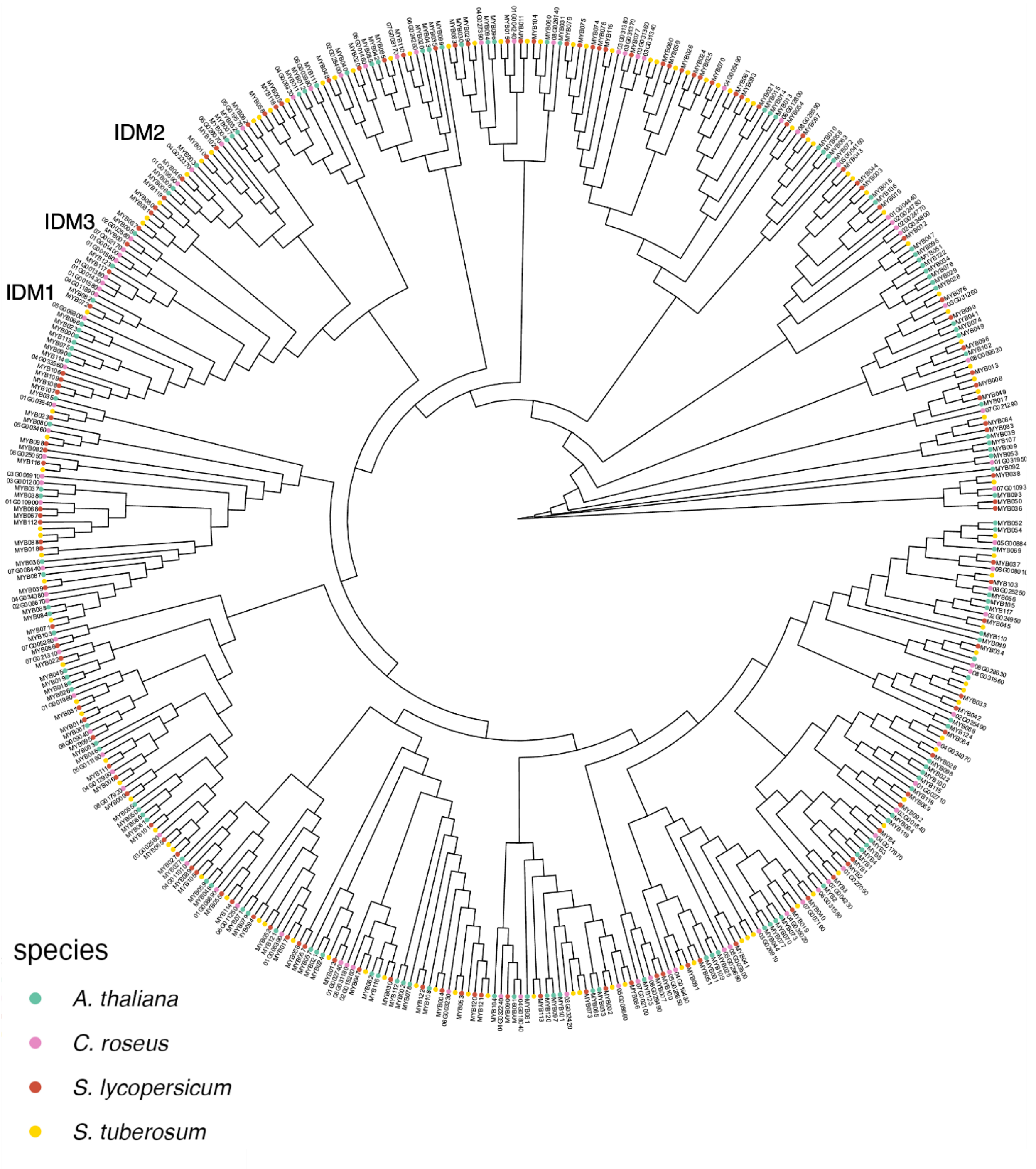


**Supplementary Fig. 8. Phylogeny of MYB domain proteins in the *C. roseus* genome.**

The colors of tips indicate species of origin. Idioblast specific MYBs are labeled.

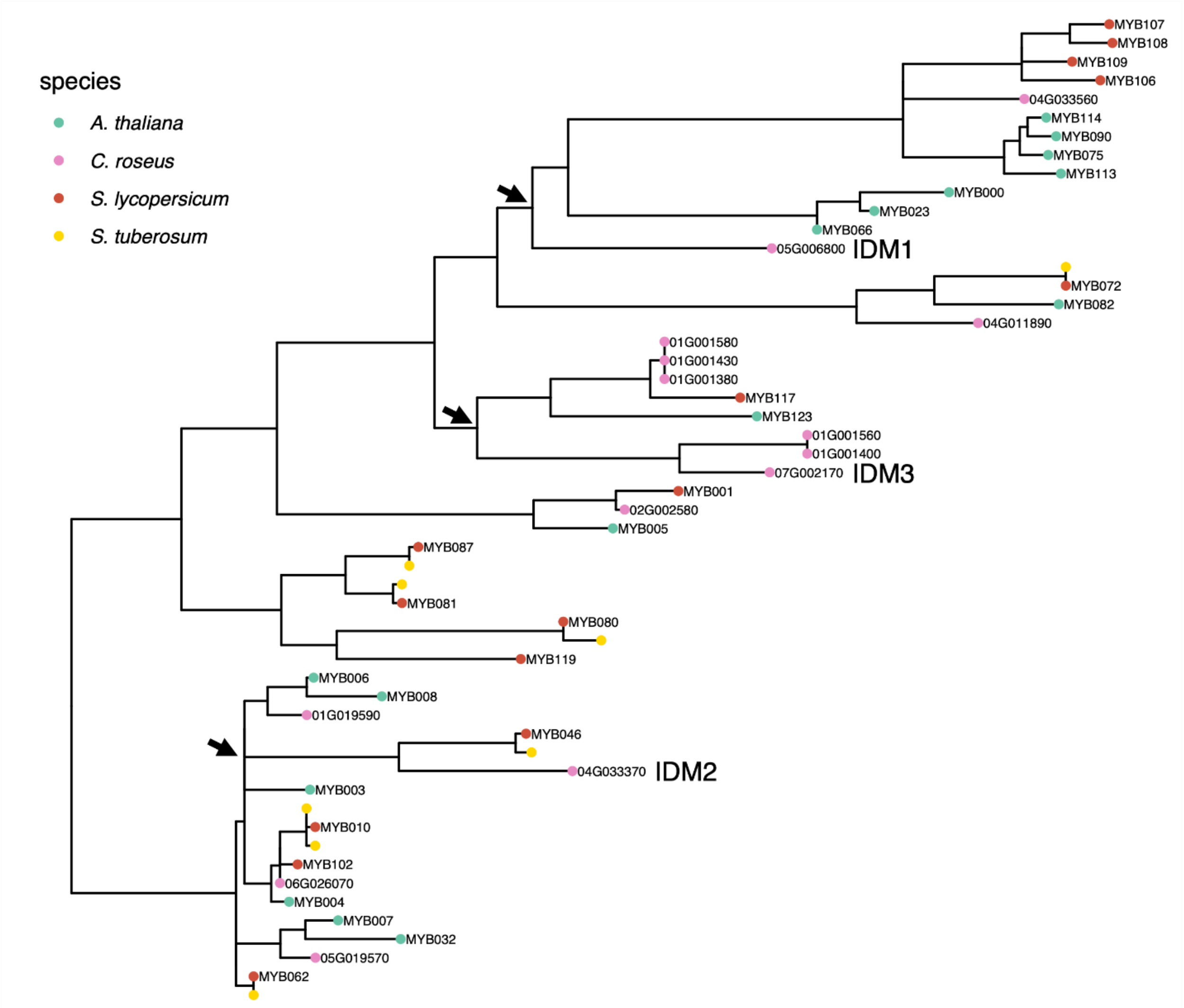


**Supplementary Fig. 9. Enlarged phylogeny containing IDM factors.**

The colors of tips indicate species of origin. Idioblast specific MYBs are labeled. Arrows indicate nodes most recently shared with an Arabidopsis MYB TF.

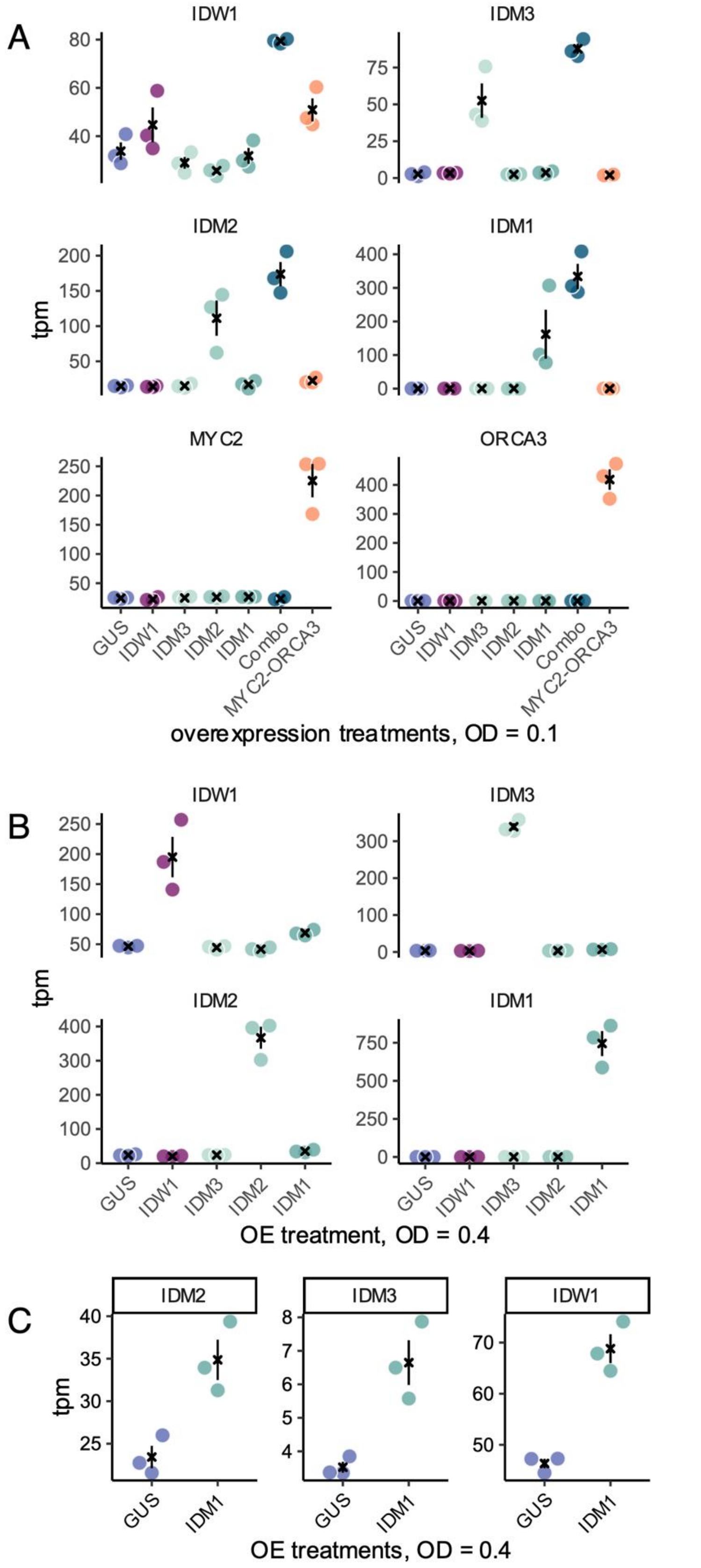

A
B
C
IDW1
IDM3
IDM2
IDM1
MYC2
ORCA3
tpm
GUS
IDW1
IDM3
IDM2
IDM1
Combo
MYC2-ORCA3
overexpression treatments, OD = 0.1
OE treatment, OD = 0.4
OE treatments, OD = 0.4

**Supplementary Fig. 10. Gain-of-function experiments on idioblast specific TF candidates using overexpression assay.**

y-axis indicates expression level, in units of transcription per million. Each dot is a biological replicate, color coded by the overexpression treatment. Error bars indicate average and standard error. Black × indicates average.

A. Mean separation plots showing the extent of TF overexpression in each infiltration treatment for the 0.1 OD experiment.

B. Mean separation plots showing the extent of TF overexpression in each infiltration treatment for the 0.4 OD experiment.

C. Mean separation plots showing the expression of IDM2, IDM3, and IDW1 in GUS and IDM1 treatments for the 0.4 OD experiment.

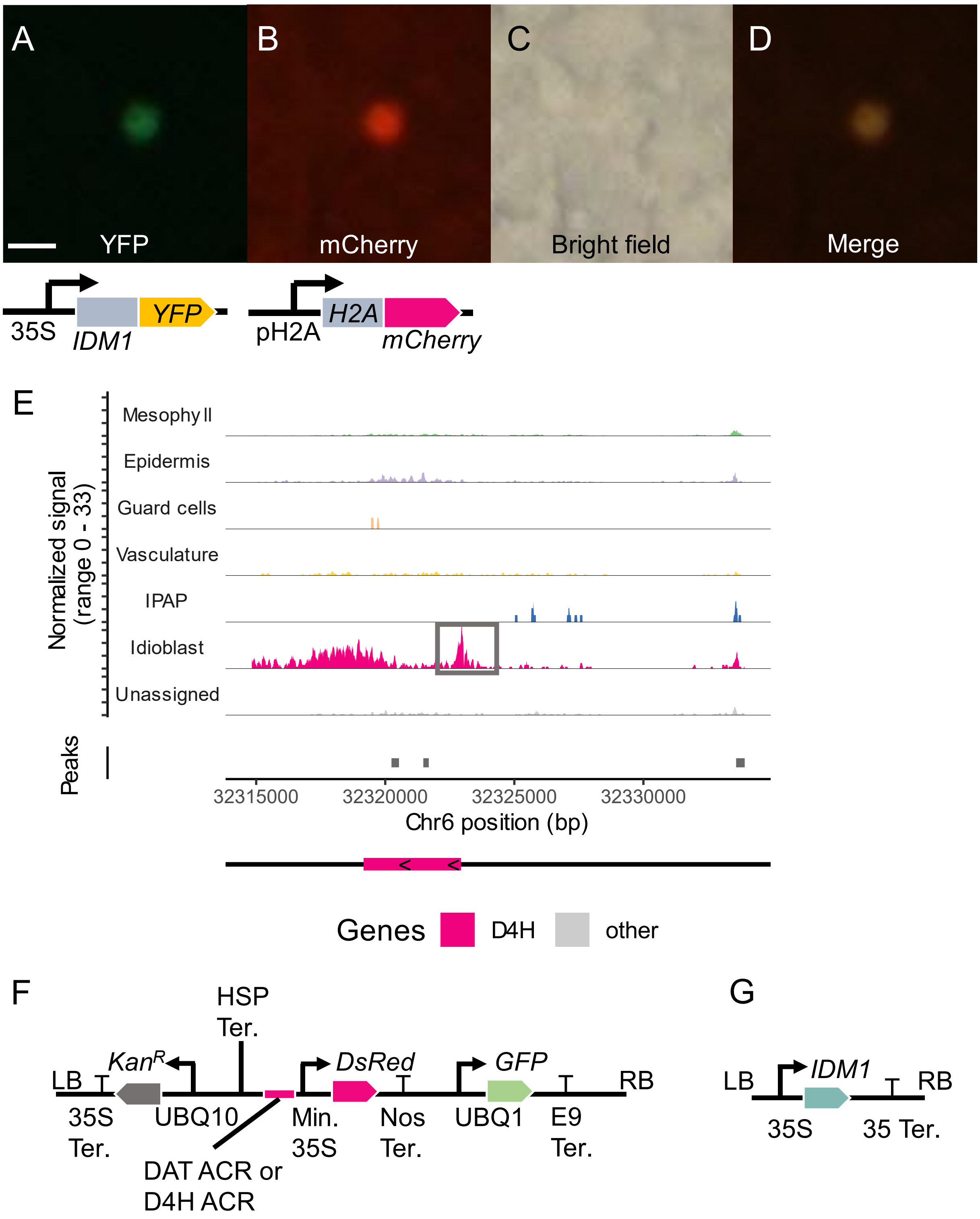


**Supplementary Fig. 11. Characterization of IDM1.**

A-D: Nuclear localization of IDM1.

E. Coverage plot showing ATAC-seq signals at the D4H locus. Box highlights the accessible chromatin region directly upstream of D4H.

F. A diagram for reporter constructs. LB. T-DNA left border. 35S Ter. 35S terminator. $Kan^R$. Kanamycin resistance gene. UBQ10: Arabidopsis UBQ10 promoter. HSP ter. Arabidopsis Heat shock protein terminator. Min. 35S. Minimal 35S promoter. Nos Ter. nopaline synthase terminator. UBQ1: Arabidopsis UBQ1 promoter. E9 ter. Terminator from the pea rbcS-E9 gene.

G. A diagram for the IDM1 overexpression construct. 35S. Full length 35S promoter.

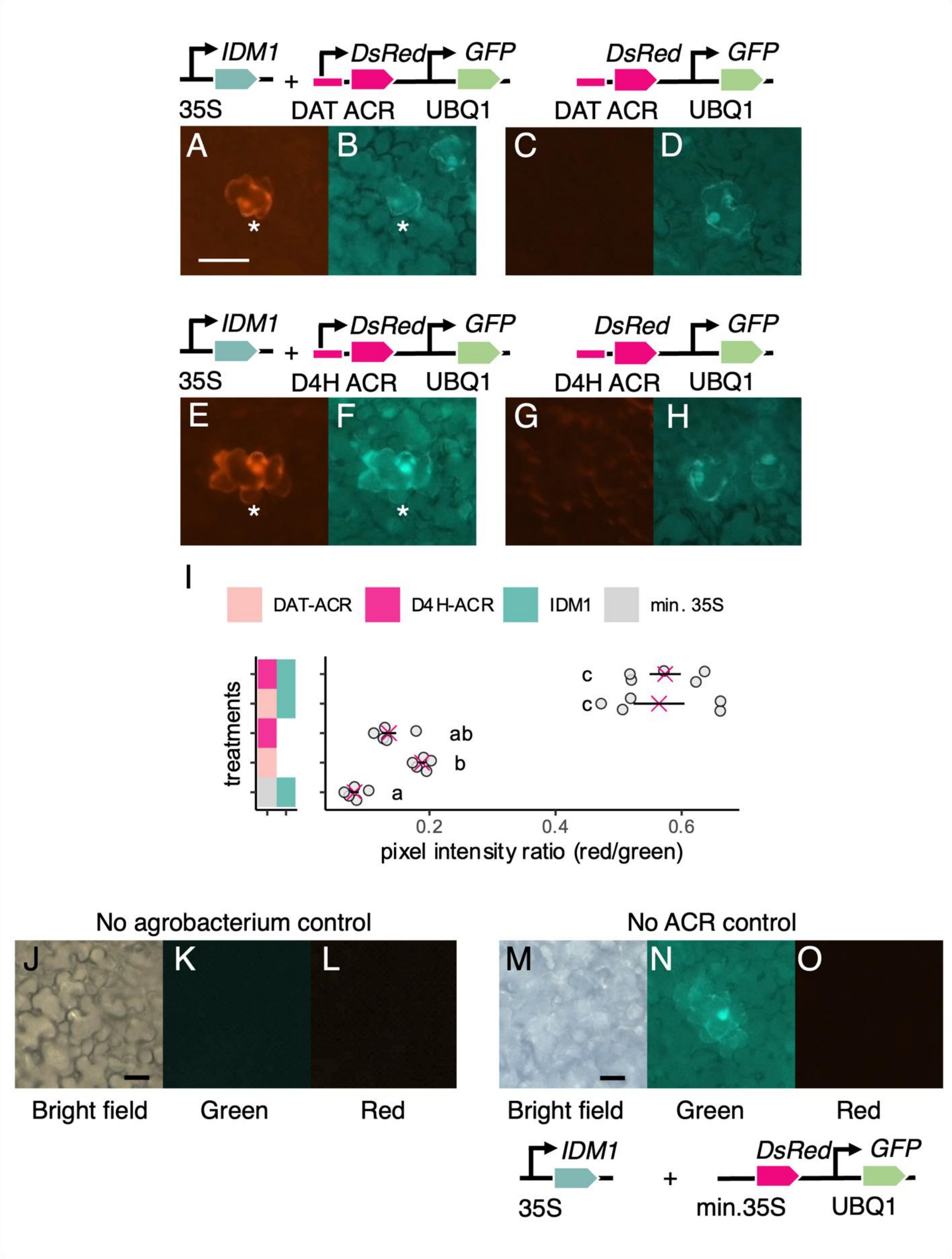


**Supplementary Fig. 12. Transactivation assays for IDM1 against DAT and D4H promoters.**

A-B: Reporter transactivation when agrobacterium strain carrying 35S:IDM1 and DAT reporter are co-infiltrated.

C-D: Control samples when only the agrobacterium strain carrying the DAT reporter is infiltrated.

E-F: Reporter transactivation when agrobacterium strain carrying 35S:IDM1 and D4H reporter are co-infiltrated.

G-H: Control samples when only the agrobacterium strain carrying the D4H reporter is infiltrated.

I. Quantification of microscope pixel intensity ratios. Color boxes on the right represent agrobacterium strains infiltrated. The first column indicates the reporter constructs (DAT, D4H, or minimal 35S promoter control). The second column indicates whether 35S:IDM1 is co-infiltrated. Error bar represents average and standard error. Pink × indicates average.

J-L: Microscope images of petals infiltrated with infiltration buffer (no agrobacterium control). No fluorescent signal can be detected in either the green or red channel.

M-O: Microscope images of petals co-infiltrated with 35S:IDM1 and a control reporter without accessible chromatin region. No fluorescent signal can be detected in the red channel.

Bar indicates 20 µm. ACR: accessible chromatin region.
